# Genome-Wide High Resolution Expression Map and Functions of Key Cell Fate Determinants Reveal the Dynamics of Crown Root Development in Rice

**DOI:** 10.1101/2020.06.11.131565

**Authors:** Tushar Garg, Zeenu Singh, Kunchapu Chennakesavulu, Anuj K. Dwivedi, Vijina Varapparambathu, Raj Suryan Singh, Khrang Khrang Khunggur Mushahary, Manoj Yadav, Debabrata Sircar, Divya Chandran, Kalika Prasad, Mukesh Jain, Shri Ram Yadav

**Affiliations:** Department of Biosciences and Bioengineering, Indian Institute of Technology, Roorkee, Uttarakhand, India; School of Computational and Integrative Sciences, Jawaharlal Nehru University, New Delhi, India; School of Biology, Indian Institute of Science Education and Research, Thiruvananthapuram, Kerala, India; Laboratory of Plant-Microbe Interactions, Regional Center for Biotechnology, NCR Biotech Science Cluster, Faridabad, Haryana, India

## Abstract

Shoot borne adventitious/crown roots (AR/CR) shape up the root architecture in grasses. Mechanisms underlying initiation and subsequent outgrowth of CR remain largely unknown. Here, we provide genome-wide modulation in the landscape of transcriptional signatures during distinct developmental stages of CR formation in highly derived grass species, rice. Our studies implicate the role of potential epigenetic modifiers, transcription factors and cell division regulators in priming the initiation of CR primordia followed by progressive activation of conserved transcription regulatory modules to ensure their outgrowth. In depth analysis of spatio-temporal expression patterns of key cell fate determinants and functional analyses of rice *WUSCHEL RELATED HOMEOBOX10* (*OsWOX10*) and *PLETHORA* (*OsPLT1*) genes reveal their unprecedented role in controlling root architecture. We further show that *OsPLT1* activates local auxin biosynthesis and forms an integral part of *ERF3-OsWOX11-OsRR2* regulatory module during CR primordia development. Interestingly, *OsPLT* genes, when expressed in the transcriptional domain of root-borne lateral root primordia of *Arabidopsis plt* mutant, rescued their outgrowth demonstrating the conserved role of *PLT* genes in root primordia outgrowth irrespective of their developmental origin. Together, these findings unveil the molecular framework of cellular reprogramming during trans-differentiation of shoot tissue to root leading to culmination of robust root architecture in grass species which got evolutionary diverged from dicots.

## INTRODUCTION

In rice (*Oryza sativa*), the mature root system is mainly composed of post-embryonic shoot-borne adventitious/crown roots (ARs/CRs) and root-borne lateral roots (LRs). The origin of various post-embryonic roots is highly diversified in plant species. For example, the pericycle cells at the xylem pole of hypocotyl give rise to ARs in *Arabidopsis*, whereas ARs/CRs are developed from the innermost ground meristem cells peripheral to the vascular cylinder at the stem base in rice (Itoh et al. 2005, Rebouillat et al. 2009, Bellini et al. 2014). Similarly, LRs originate from the xylem pole pericycle cells of *Arabidopsis* primary root (PR), whereas rice LRs originate from endodermal and pericycle cells located opposite to the protophloem of PR and CRs (Itoh et al. 2005, Rebouillat et al. 2009; Lavenus et al. 2013; Bellini et al. 2014). Despite gross morphological similarities among various root types in cereals, diverged root-type and species-specific regulatory mechanisms also exist (Kitomi et al. 2011a, Orman-Ligeza et al. 2013; Meng et al., 2019).

The establishment of founder cells for rice crown root primordia (CRP) requires an induction phase for cell cycle reactivation in a localized domain of the innermost ground tissues of stem base to produce initial cells for CRP (Itoh et al., 2005; Guan et al., 2015). These initial cells, originated from shoot tissues, then divide and their daughter cells trans-differentiate to produce root tissues. Different developmental events of *de novo* CR organogenesis have been divided into seven stages, starting from the initial cell establishment for CRP until their emergence (Itoh et al., 2005). Still, only a handful of CRP expressed genes are identified and global gene architecture during CRP development is not explored. *ADVENTITIOUS ROOTLESS 1* (*ARL1*)/*CROWN ROOTLESS 1* (*OsCRL1*) is amongst the early regulators of rice CRP development as in the *crl1* mutants early cell division is suppressed (Inukai et al., 2001; 2005; Liu et al. 2005). Other TFs, such as *ETHYLENE-RESPONSIVE FACTOR3* (*OsERF3*), *OsTOC168/OsAP2/ERF-40, CROWN ROOTLESS5* (*OsCRL5*), and *WUSCHEL-RELATED HOMEOBOX 11* (*OsWOX11*), *CYTOKININ RESPONSE REGULATOR 2* (*OsRR2*) and *SQUAMOSA PROMOTER BINDING PROTEIN-LIKE3* (*OsSPL3*) play key roles at different stages of CR development in rice (Kitomi et al. 2011a; 2011b, Zhao et al. 2009, 2015, Neogy et. al., 2019, Shao et. al., 2019).

Auxin maxima activates the signaling that initiate program for root founder cell specification and also ensures entire post-embryonic root development (De Rybel et al. 2010, Yadav et al. 2010, Lavenus et al. 2013). Mutation or over-expression of rice genes regulating local auxin biosynthesis, distribution, and signaling, such as *YUCCA* genes, *CROWN-ROOTLESS4/OsGNOM1* (*OsCRL4/OsGNOM1*) and *PINFORMED1* (*OsPIN1*) display defects in CR development (Yamamoto et al., 2007; Kitomi et al. 2008, Liu et al. 2009, Li et al., 2019). Auxin signaling activates expression of TFs, which in turn, regulate and integrate the signaling pathway during CR organogenesis (Inukai et al. 2005, Liu et al. 2005, Kitomi et al. 2011b, Zhao et al. 2009, 2015, Coudert et al. 2015, Zhang et al., 2018, Lavarenne et al., 2019, Neogy et al., 2019; Mao et al., 2020).

In this study, we investigated the global gene network operational during CRP specification and differentiation in rice. The genome-wide laser capture microdissection coupled with RNA sequencing (LCM-seq) based transcriptome analysis revealed a genetic blueprint for the localized developmental reprogramming during the trans-differentiation process and demonstrated spatio-temporal regulated expression patterns of a set of transcription factors during CRP initiation and outgrowth. The detailed functional analysis of stem cell promoting factors, *OsWOX10* and *OsPLT1* genes, reveal their previously unrecognized role in controlling root architecture. *OsWOX10* promotes adventitious root development across the species and has a non-redundant function in regulating rice CR development *via OsRR2. OsPLT1* function is necessary and sufficient to promote CR and LR development in rice. Strikingly, *OsPLT1* stimulates local auxin biosynthesis by directly activating *OsYUC1* and *OsYUC3* and also activates the expression of *ERF3, OsWOX11* and *OsRR2* genes in this process. Interestingly, *OsPLT1* can rescue lateral root primordia outgrowth defects in *Arabidopsis plt* triple mutants. Taken together, our studies reveal a novel regulatory network operating during rice CR development and uncover a key function and putative mechanisms of *OsWOX10* and *OsPLT1* in instructing the root architecture in rice.

## RESULTS

### Laser Capture Microdissection and Global Gene Expression Profile During CRP Initiation and Outgrowth

To dissect the determinants of cell fate specification and generate high-resolution temporal gene expression map of rice CRP at their progressive developmental stages, we performed LCM coupled with RNA sequencing (LCM-seq) to profile transcripts from CRP during their initiation and outgrowth. During the initiation stage, cell identity of CRP initials is established, which eventually produce initials for epidermis-endodermis, root cap, and central stele (**Figure 1A-1C**). Subsequently, CRP progress to outgrowth stage, where tissue patterning leads to the fundamental tissue organization (**Figure 1A and 1D**). Tissues from eleven initiation-stage (**Figure 1E and 1E’**) and ten outgrowth-stage CRP were collected (**Figure 1F and F’**). The innermost ground tissues peripheral to the vascular cylinder have the competence to initiate a CR-specific developmental program when proper cues are perceived, therefore these tissues were collected as a control (**Figure 1G and 1G’**). Total RNAs extracted from the LCM collected cells were subjected to RNA sequencing and expression profiles of all the rice genes were revealed. In order to uncover common patterns of gene expression, fuzzy c-means clustering was performed on the transcriptomic data, as studied earlier (Harrop et al., 2016) and eight clear clusters with distinct expression patterns were observed (**Figure 1H**). GO enrichment analysis of these clusters provided association of expression patterns with biological processes. Clusters 1 and 2 genes, expressed in both the stages were mainly enriched with GO terms related to DNA replication and gene expression, cluster 3 genes, specifically induced during CRP initiation, were associated with cell cycle and division processes, whereas hormonal signaling genes were enriched among cluster 4 genes which are induced during CRP outgrowth (**Figure 1H, 1I**), indicating an order of events to set developmental program. However, cluster 5-8 genes, which are mostly associated with stimuli responses and other developmental processes, were repressed during CRP development (**Figure 1H, 1I**).

**Figure 1:**
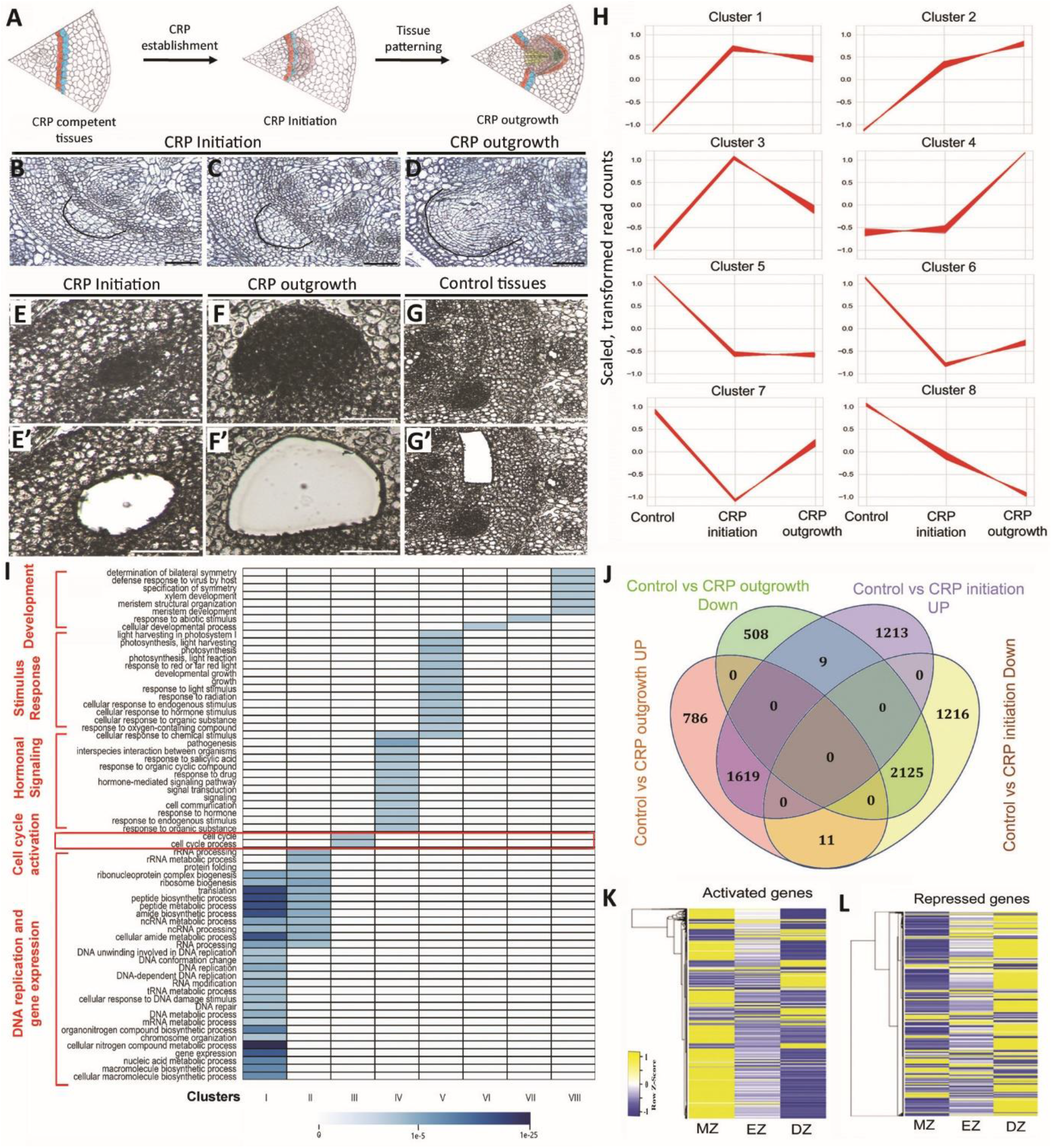
Laser capture microdissection-RNA sequencing (LCM-seq) analysis of developing crown root primordia (CRP). (A) Schematic diagram showing CRP competent tissues and developing CRP at the stage of initiation and outgrowth. (B-C) Cross sections of rice stem base with CRP at initiation (B, C) and outgrowth stage (D). (E-G’) LCM of CRP during their initiation (E, E’) and outgrowth (F, F’). Innermost ground tissues prior to CRP initiation were collected as control (G, G’). (E)-(G) are before and (E’)-(G’) after LCM. Eleven CRP at the stage of initiation and ten outgrowing CRP were collected for RNA-seq analysis. (H) Patterns of gene expression during CRP initiation and outgrowth. (I) Heat-map showing mostly distinct GO terms enrichment in different clusters. Log2 fold change ≥ 1 with p<0.05 parameter was considered, and p-value was used to generate the heatmap. (J) Venn diagram showing common and unique differentially expressed genes (DEGs) during CRP initiation and outgrowth. (K, L) Expression pattern of DEGs in different zones of emerged roots. (MZ, meristematic zone; EZ, elongation zone; DZ, differentiation zone). Bars= 50µm in (B)-(G’).

The differential gene expression analysis revealed that expression of 2429 genes (1213 induced and 1216 repressed) during CRP initiation, and 1294 genes (786 up-regulated and 508 down-regulated) during CRP outgrowth, was specifically de-regulated, as compared to control tissues (**Figure 1J; Supplemental Dataset 1-2**). However, a total of 3744 (1619 induced and 2125 repressed) genes were commonly regulated during CRP initiation and outgrowth stages (**Figure 1J; Supplemental Dataset 2**). The expression of 1035 genes was higher at the initiation-stage CRP as compared to outgrowing CRP, whereas an opposite pattern was observed for 1333 genes (**Supplemental Dataset 1**). We further observed that the activated genes during CRP development showed a higher expression in the actively dividing meristematic zone than the differentiation zone, whereas the CRP repressed genes showed opposite pattern between meristematic and differentiation zone of emerged roots (**Figure 1K and 1L**). Thus, our data show that a dynamic transcriptional reprogramming is instrumental for the cell fate change, cell division and differentiation.

### Specific Induction of Cell Cycle Genes, and Epigenetic Modifiers During CRP Initiation

Next, GO enrichment analysis of the differentially expressed genes (DEGs) provided deeper insights into the various biological processes associated during CRP development. The initiating CRP were exclusively enriched with genes regulating hormonal levels, transcription pre-initiation, RNA processing, cell cycle, and organ development, whereas GO terms related to metabolic processes are associated with the genes exclusively induced during CRP outgrowth (**Supplemental Figure 1A**). Other set of genes which are associated with biological processes such as, hormonal metabolism and signaling, gene regulation, nucleic acid metabolism, cell division, and organ development including post-embryonic root organogenesis displayed higher expression in the initiating CRP and their expression reduced in the outgrowing CRP (**Supplemental Figure 1B**). Collectively, this suggests that primary biological and regulatory processes required for establishment and differentiation of CRP are associated with genes highly expressed during CRP initiation, whereas metabolic processes were enriched in outgrowing CRP. Further, the geneset enrichment analysis (GSEA) of transcriptional regulators showed that several members of PHD, SWI/SNF, SET, GNAT, and Jumonji gene families, involved in epigenetic and chromatin-remodelling-mediated pre-transcriptional gene regulation, are largely induced during CRP initiation (**Figure 2A; Supplemental Table 1**). To validate involvement of putative epigenetic regulators in CRP initiation, we studied temporal and spatial expression pattern of a few chromatin-remodelling genes; two PHD-domain containing factors from trithorax group proteins (*OsTRX1, Os09g04890* and *OsATXR6, Os01g73460*) and a SWIB/MDM2 domain containing protein, *Os03g55310*, during CRP establishment and differentiation. RNA *in situ* hybridization showed that onset of their activation was at very early stage in the CRP founder cells **(Figure 2B, 2D and 2F)** and they continue to express during CRP outgrowth **(Figure 2C, 2E and 2G; Supplemental Figure 2A-2C)**, corroborating their putative function during CRP specification and differentiation.

**Figure 2:**
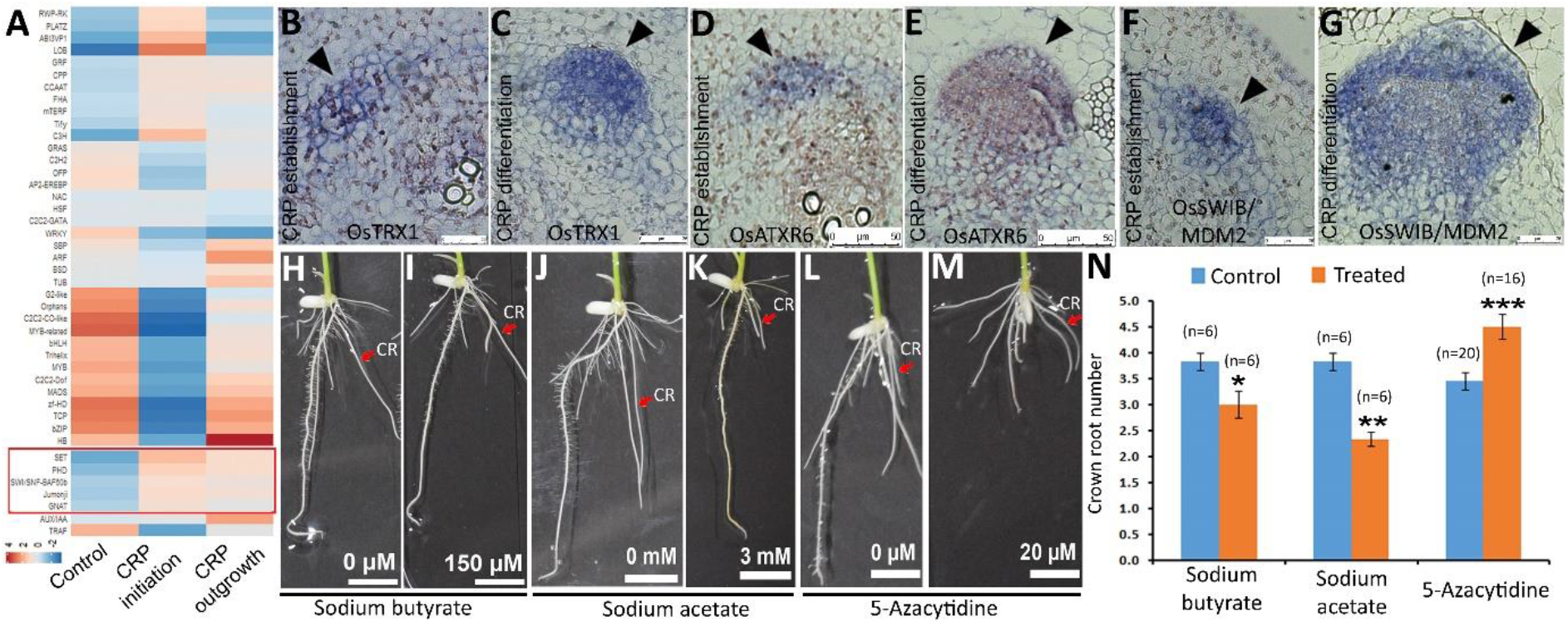
Epigenetic regulation is involved in controlling rice root architecture (A) Geneset enrichment analysis of TFs. Gene families in red box are involved in epigenetic regulation. (B-G) Spatial expression pattern analysis of three putative genes involved epigenetic regulation. (H-K) Alteration in histone acetylation by treating with 150µM sodium butyrate (H, I) or 3mM sodium acetate (J, K) results in altered rice root architecture with reduced CR number. (L, M) Inhibition of DNA methylation by treating with low doses of 5-azacitidine (20µM) increases CR number. (N) Quantitative representation of CR numbers upon treatments with these drugs. (*p≤0.05; **p≤0.005; ***p≤0.001; two-sample t-test). Site of CRP establishment and developing CRP are highlighted with black arrowhead in (B)-(G). Red arrows mark CR in (H)-(M). (CRP, crown root primordia; CR, crown root). Bars= 25µm in (B), (C), (F), (G), 50µm in (D), (E), 1cm in (H)-(M).

Further, to functionally support involvement of epigenetic regulation during CRP development, we studied consequence of interference of epigenetic modifiers on rice root architecture. Sodium butyrate (inhibits histone deacetylase) and sodium acetate (acetylates histones) are known to promote histone acetylation, whereas 5-azacitidine is a potent inhibitor of DNA methylation (Qiu et al., 2019; Zhang et al., 2013). We observed that overall root architecture was altered upon treatments with these pharmacological drugs **(Figure 2H-2M)**. The CR number was reduced upon treatment with sodium butyrate and sodium acetate whereas it was increased upon 5-azacitidine treatment **(Figure 2N)**. Corroborating with these observations, we also observed that expression level of CRP expressed transcription factors, *OsWOX10* and *OsPLT1* was altered upon sodium acetate and 5-azacitidine treatments in an opposite manner (**Supplemental Figure 2D**). Thus, these data together indicate involvement of histone acetylation and DNA methylation-dependent epigenetic regulation during CRP development.

### Progressive and Spatial Activation of Auxin Signaling During CRP Development

Coordinated auxin biosynthesis, homeostasis and distribution is required for generating local auxin maxima which is an early event to activate auxin signaling in the localized domains to trigger post-embryonic root developmental programs (Benková et al., 2003; Dubrovsky et al., 2008; Yadav et al., 2011). We observed dynamic and progressive activation of auxin signaling during CRP development. An auxin biosynthesis gene of the YUCCA gene family, *OsYUC3* is strongly expressed in the competent tissues prior to CRP initiation (**Supplemental Figure 3A; Supplemental Dataset 3**). This was followed by robust activation of four YUCCA genes (*OsYUC1, 4, 7*, and *9*) during CRP initiation. Notably, the expression of all except *OsYUC1* was progressively increased, as CRP progress from initiation to outgrowth stage (**Supplemental Figure 3A; Supplemental Dataset 3**). However, the GH3 genes, which reduce active pool to auxin to maintain auxin homeostasis, displayed opposite pattern. The expression of five GH3 genes (*OsGH3*.*1, OsGH3*.*2, OsGH3*.*4, OsGH3*.*8*, and *OsGH3*.*13*) was activated during CRP initiation, but the expression of all except *OsGH3*.*2* was progressively reduced in outgrowing CRP (**Supplemental Figure 3A; Supplemental Dataset 3;** Singh et al., 2021).

Auxin signaling is activated by de-repressing auxin response factors (ARF) through auxin-dependent proteolytic degradation of Aux/IAA proteins, a class of negative regulators of auxin signaling. Consistent with the above observations, we also noticed a progressive transcriptional repression of Aux/IAA genes during CRP progression. The expression of 5 *OsIAA* genes was specifically induced and only *OsIAA13* is repressed during CRP initiation, whereas 4 *OsIAA* genes were repressed and none was exclusively induced in outgrowing CRP (**Supplemental Figure 3B; Supplemental Dataset 3**). Furthermore, the expression levels of 11 *OsIAA* genes were repressed but only 3 genes were induced when CRP progress from initiation to outgrowth stage (**Supplemental Dataset 3**). Further, a sharp transcriptional activation of *OsARF16* and *OsARF10* was observed during CRP initiation (**Supplemental Figure 3C; Supplemental Dataset 3**). While the expression level of *OsARF16* was declined, *OsARF10* transcript level was maintained during CRP outgrowth. *OsARF8* and *OsARF75* were exclusively induced in the outgrowing CRP. The transcription of *OsARF8* and *OsARF22* was progressively induced and the expression of a set of *OsARFs* was decreased in the developing CRP (**Supplemental Figure 3C; Supplemental Dataset 3**). These observations suggest that a controlled and progressive activation of auxin biosynthesis, and signaling genes might generate a spatio-temporal auxin signaling modules required for CRP initiation and outgrowth.

To further confirm this observation, we studied the spatio-temporal activation of auxin signaling by monitoring auxin response during rice CRP development. Cross sections of stem base containing developing CRP of transgenic rice lines expressing DR5-erYFP construct was hybridized with anti-sense RNA probes and anti-GFP antibodies. RNA *in situ* hybridization and immunohistochemistry analysis revealed that auxin response is initiated in a localized domain at very early stage of CRP specification and no signal was detected in CRP of wild-type plant (**Figure 3A and 3A1; Supplemental Figure 3D**). During later stages of CRP outgrowth, auxin signaling is more at the tip of CRP (**Figure 3A 2-3C**), which eventually gets restricted to QC, columella and initial cells of the emerged and growing root tip (Yang et al., 2017). All these observations together suggest that auxin signaling is activated at the onset of CRP program initiation and eventually culminate a robust auxin signaling during CRP outgrowth.

**Figure 3:**
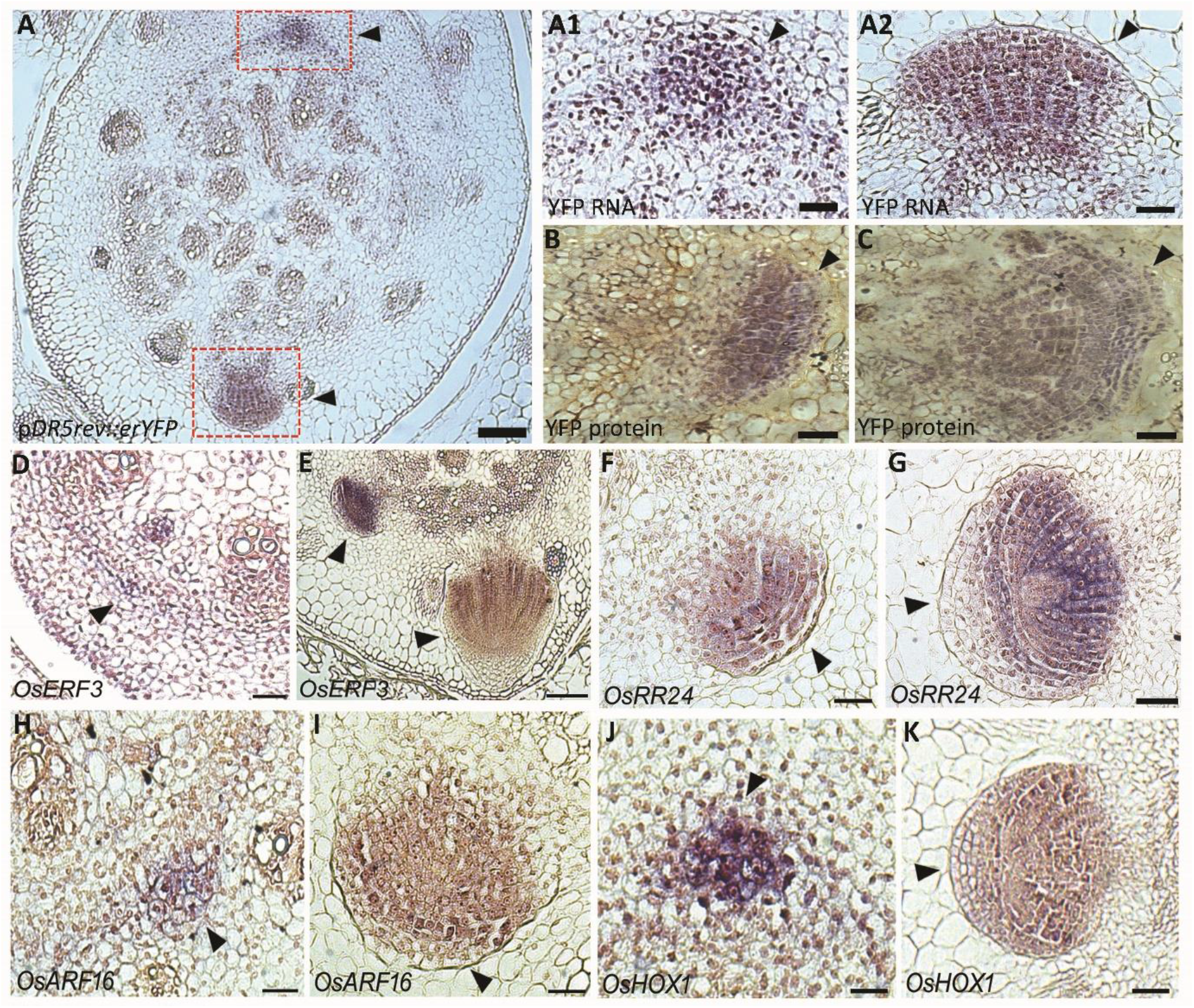
Auxin response and tempo-spatial expression pattern of selected TFs during CRP development. (A-C) Spatially regulated auxin response analysed by pDR5rev::erYFP expression pattern during CRP development in rice. RNA *in situ* hybridization using anti-sense RNA probes (A-A2) and immuno-histochemistry using anti-GFP antibody (B, C) were performed. CRP marked in red box in (A) are enlarged in (A1) and (A2). (D-L) RNA *in situ* hybridization of four selected TFs. (D, E) AP2-domain containing *OsERF3*, a regulator of CRP initiation. (F, G) Type-B cytokinin response regulator, *OsRR24*. (H, I) Auxin response factor, *OsARF16*. (J, K) Homeobox containing TF, *OsHOX1*. Developing CRPs probed with DIG-labeled anti-sense RNA probes of respective genes. Site of CRP and developing CRP are highlighted with black arrowhead. (CRP, crown root primordia). Bars= 25 µm in (A1)-(G), and (J) and 100 µm in (A), (I), and (K).

### Stage-specific Signatures of Transcriptional Regulators During CRP Development

Transcription factors (TFs) are master regulators of cell fate determination. We observed that the expression of 191 TFs (83 induced and 108 repressed) was exclusively de-regulated during CRP initiation, whereas 96 TFs (41 up-regulated and 55 down-regulated) were exclusively de-regulated during CRP outgrowth (**Supplemental Figure 4A; Supplemental Dataset 4**). Positioning and maintenance of stem cell niche (SCN) close the QC cells in the root meristem is essential for CRP growth. This is regulated by gene regulatory modules involving TFs, *SHORT ROOT* (*SHR*), *SCARECROW* (*SCR*), *PLETHORA* (PLTs) and *WOX5* (Sabatini et al., 2003; Aida et al., 2004; Dinneny and Benfey, 2008). LCM-seq data showed that both rice paralogous *SHR* (*OsSHR1, OsSHR2*), and *SCR* genes (*OsSCR1* and *OsSCR2*), along with *OsPLTs* (*OsPLT1-OsPLT6*), and *WOX5* homolog (*QHB*) displayed strong and dynamic expression during CRP development (**Supplemental Figure 4B, 4C; Supplemental Table 2**), suggesting that a mechanism with conserved regulators is involved in establishing functional SCN in the developing CRP.

The expression of many TFs, such as *OsCRL1, OsCRL5, OsERF3* and *OsRR2* that promote CRP initiation, is regulated by auxin signaling (Inukai et al., 2005; Kitomi et al., 2011a; Zhao et al., 2015). We observed that out of 83 CRP initiation-specific induced TFs, 15 TFs that include *OsCRL1, OsERF3, OsWOX10, OsHOX12, OsMFS1, OsHOX1, OsSta2* were auxin inducible, whereas remaining including *FREEZY PANICLE* (*FZP*), CRL1-like *OsCRL1L2, OsLBD37/ASL39, QHB, ROC4, OsRR24* and *OsHB34* were auxin non-responsive (**Supplemental Figure 5A, 5A1; Supplemental Dataset 5**). Similarly, 6 of 41 TFs, specifically induced during CRP outgrowth, involving *OsAP2/EREBP127, OsNAC10, HSFB4A* and *OsRR5*, were auxin inducible (**Supplemental Figure 5B, 5B1; Supplemental Dataset 5**). However, 16 of 112 TFs whose expression was activated during both stages, including *OsDH1, OsNAC039, OsERF61, OsbHLH1, bZIP78*, and *OsNAC139*, are targets of auxin signaling, whereas remaining TFs like *OsGATA10, 15, OsLBD1-8, OsGRFs* and *OsMADS17* are not regulated by auxin (**Supplemental Figure 5C, 5C1; Supplemental Dataset 5**). Our qRT-PCR analysis validated auxin induction of a few selected CRP expressed TFs upon IAA treatment (**Supplemental Figure 6A-6D**). Thus, the study identified stage-specific, auxin-dependent and auxin-independent transcriptional regulators of CRP initiation and outgrowth.

### Spatio-Temporal Expression Pattern of Transcription Factors in Developing CRP

Next, we studied the detailed temporal and spatial expression of a few selected genes exclusively activated during CRP initiation in LCM-seq data to uncover the onset and dynamic expression pattern during CRP development. We selected an AP2 domain-containing transcription factor (*OsERF3*), an auxin response factor (*OsARF16*), a cytokinin response regulator (*OsRR24*), and an auxin-responsive homeobox-containing transcription factor (*OsHOX1*) for RNA *in situ* hybridization using DIG-UTP labeled anti-sense and sense RNA probes. We observed that all of these genes were specifically and strongly expressed in developing CRP, and other tissues did not show any expression above the background level (**Figure 3D-3K**). The onset of expression of *OsERF3, OsARF16*, and *OsHOX1* was in the localized domains of tissues peripheral to the vascular tissues during CRP establishment (**Figure 3D, 3H and 3J**) and they continue to express in the developing CRP (**Figure 3E, 3I, 3K**). The expression of *OsRR24* was detected at slightly later stages when CRP is already established (**Figure 3F**). *OsERF3* and *OsRR24* were expressed throughout the early CRP, but in the outgrowing CRP, the expression is reduced in the root cap tissues (**Figure 3D-3G**). The expression of *OsARF16* was initiated at the early stage of CRP specification and continued to express in the differentiating CRP. The expression was more restricted towards the apical region of the CRP as compared to the base of the CRP (**Figure 3H and 3I**). In the outgrowing CRP, the expression of *OsRR24* was reduced in the QC and surrounding initials of the ground and vascular tissues, and their immediate daughter cells (**Figure 3G**). The *OsHOX1* showed a very strong expression at the site of CRP specification and relatively uniform expression in the late CRP (**Figure 3J and 3K**). However, cross-sections hybridized with sense probes did not show any signal above the background level (**Supplemental Figure 7A-7D**). These results confirm that the expression of these TFs is confined to developing CR primordia and suggests a strict necessity of their spatial regulation during CR development.

### *OsWOX10* Promotes Adventitious Root Formation

*Arabidopsis WOX11* and *WOX12*, members of WOX gene family, regulate first-step cell fate transition during root organogenesis (Lian et al., 2014; Liu et al. 2014; Hu and Xu, 2016). Of three rice members of *WOX11/12* clade, we studied function of a related rice WOX gene **(Figure 4A**), *OsWOX10*, whose expression was sharply activated during CRP initiation **(Supplemental Figure 6B)**. Importantly, the expression of *OsWOX10* is strongly induced by auxin signaling (**Supplemental Figure 6D;** Neogy et al., 2019). Our detailed temporal and spatial expression pattern analysis of *OsWOX10* transcript localization demonstrated that *OsWOX10* transcription is specifically activated in the founder cells of CRP, prior to their establishment that also coincide with auxin maxima (**Figure 4B**) and continue to express in the initiating and outgrowing CRP (**Figure 4C and 4D**). These observations suggest that auxin signaling activates *OsWOX10* expression at the onset of CRP specification.

**Figure 4:**
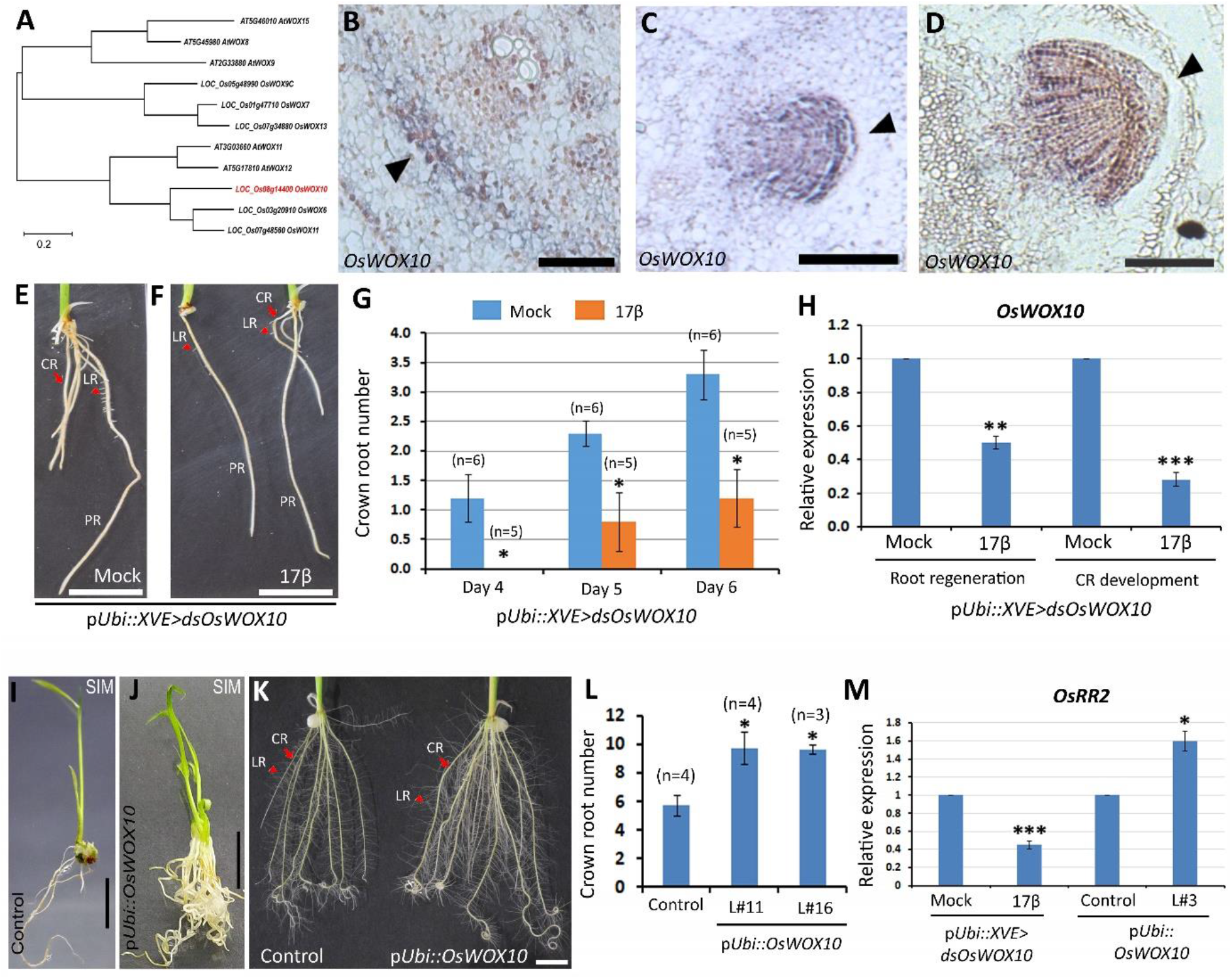
*OsWOX10* promotes adventitious root development. (A) Phylogenetic analysis of WOX members of intermediate clade show that *OsWOX10* is closely related to *Arabidopsis WOX11/12*. (B-D) Onset and tissue-specific expression pattern of *OsWOX10* during rice CRP development. *OsWOX10* expression is initiated in the CR founder cell (B) and later expression is specific to developing CRP (C, D). (E, F) CR number is reduced when *OsWOX10* is down-regulated upon 17β-estradiol treatment (F) in pUbi::*XVE>dsOsWOX10* lines. (G) Quantitative representation of crown root number (*p≤0.05; two-sample t-test). (H) Partial down-regulation *OsWOX10* upon 17β-estradiol treatment during root regeneration in T0 and CR development in T1 generation (**p≤0.005; ***p≤0.001; two-sample t-test). (I-K) Consequence of ectopic over-expression of *OsWOX10*. Extensive rooting is seen in shoot induction (J) media as compared to control (I) during plantlet regeneration. (K, L) CR number was also increased upon *OsWOX10* over-expression in T1 generation (*p≤0.05; two-sample t-test). (M) *OsWOX10* activates expression of *OsRR2*, as revealed by qRT-PCR analysis (*p≤0.05; ***p≤0.001; two-sample t-test). Site of CRP and developing CRP are highlighted with black arrowhead in (B)-(D). Red arrows and arrow heads mark CR and LR, respectively in (E), (F) and (K). (CRP, crown root primordia; PR, primary root; CR, crown root; LR, lateral root). Bars= 5 µm in (B), 100 µm in (C), 25 µm in (D), 1 cm in (E), (F), and (I)-(K).

To investigate the function of *OsWOX10* during CR development, an inverted repeat RNA-interference (RNAi) construct for *OsWOX10* was expressed in transgenic rice under maize ubiquitin promoter using estradiol-inducible XVE system (p*Ubi::XVE:dsOsWOX10*). Estradiol-induced down-regulation of *OsWOX10* resulted in altered root architecture and no visible effect was observed in wild-type plants (**Supplemental Figure 8A and 8B**). Lesser number of ARs were formed upon *OsWOX10* knock-down during plantlet regeneration on the root induction media (**Supplemental Figure 9A**). During vegetative development, both CR number and length was reduced in knock-down lines (**Figure 4E-4G; Supplemental Figure 9B-9F**). In these lines, the expression of *OsWOX10* was down-regulated but no significant change was observed for related WOX genes, *OsWOX11* and *OsWOX12* (**Figure 4H; Supplemental Figure 9G)**. In contrast, ectopic over-expression of *OsWOX10* under maize ubiquitin promoter (*pUbi::OsWOX10*) in rice caused extensive root formation during plantlet regeneration and vegetative development (**Figure 4I-4L; Supplemental Figure 10A-10H**). In fact, regenerated rice plantlets initiated robust rooting even in the shoot induction media (**Figure 4J**) and also developed higher number of roots in the root induction media (**Supplemental Figure 10B**). Similarly, an opposite phenotype was observed during CR development where both CR number and length was increased in over-expression lines in T1 generation (**Figure 4K-4L; Supplemental Figure 10D-10H)**. A strong over-expression of *OsWOX10* was confirmed by qRT-PCR analysis (**Supplemental Figure 10I, 10J**). These observations suggest that *OsWOX10* is required and sufficient for initiation and growth of ARs during plantlet regeneration, and CRs during vegetative phase of rice plants.

### *OsWOX10* is an Upstream Regulator of *OsRR2* during CRP Development and its Function is Conserved in *Arabidopsis*

The functional analysis of *OsWOX10* suggested that it regulates CRP initiation and also ensures their outgrowth in rice. Next, we investigated if *OsWOX10* regulates expression of known *ERF3-OsWOX11-OsRR2* regulatory module during CRP initiation and outgrowth. We did not observe a strong effect on *OsWOX11* expression in *OsWOX10* down-regulated and over-expression lines whereas the expression of *OsRR2* was significantly reduced in *OsWOX10* knock-down and increased in *OsWOX10* over-expression lines **(Figure 4M; Supplemental Figure 9H)**, suggesting that *OsWOX10* functions upstream of *OsRR2*.

*Arabidopsis WOX11* and *12* belong to same clade as of *OsWOX10* and regulate *de novo* adventitious root organogenesis by regulating expression of *LATERAL ORGAN BOUNDARIES DOMAIN29* (*LBD29*) (Liu et al., 2014). *OsCRL1* is a putative homolog of *LBD29* which regulate rice CR development (Inukai et al., 2005), suggesting conservation of regulatory mechanism during adventitious root development between rice and *Arabidopsis*. We, therefore, next studied if molecular function of *OsWOX10* is also conserved across plant species. When *OsWOX10* was ectopically over-expressed in *Arabidopsis*, using *Arabidopsis Ubiquitin 10* promoter (Siligato et al., 2016), in wild-type as well in *wox11* and *wox12* single and double mutants where endogenous *WOX11/12* genes are not functional, it was sufficient to induce a large number of ARs from the root-hypocotyl junction (**Supplemental Figure 11A and 11B**). These data suggest conserved role of *OsWOX10* in promoting AR formation, both in dicot and monocot plant species.

### Rice *PLETHORA1* is Required and Sufficient for Post-Embryonic Root Development

*PLETHORA* (*PLT*) genes represent members of AP2-domain transcription factors, that are among key cell fate determinants of root growth and development. Rice genome encodes 10 PLT genes (Li and Xue, 2011) but their function remains unknown. Our LCM-seq data reveal that the expression of six PLT genes (*OsPLT1-OsPLT6*) is induced during CRP initiation and outgrowth (**Figure 5A; Supplemental Table 2**). We functionally studied *OsPLT1* during root development. Our temporal and spatial expression pattern analysis of *OsPLT1* showed that transcription of *OsPLT1* was activated very early at the onset of CRP specification (**Figure 5B**), with continued expression throughout the CRP during their development (**Figure 5C and 5D**). Next, to uncover function of *OsPLT1* during rice CR development, loss- and gain-of-function transgenic rice lines were generated for *OsPLT1*. For down-regulation, an RNAi construct was generated using gene-specific fragment of *OsPLT1* and was expressed under maize ubiquitin promoter (p*Ubi::dsOsPLT1*). In the transgenic rice lines where *OsPLT1* was down-regulated, the root architecture was compromised (**Figure 5F**), as compared to wild-type plants (**Figure 5E)**. In these down-regulation lines, length of all root types (i.e. PR, CR and LR), was reduced (**Figure 5F; Supplemental Figure 12A-12H**). Further, number of CRP, emerged CRs and LRs was also reduced in *OsPLT1* knock-down lines (**Figure 5G-5I; Supplemental Figure 12C, 12D**). The extent of strong down-regulation of *OsPLT1* is confirmed by qRT-PCR, however the expression level of other related rice *PLT* genes was not significantly affected (**Supplemental Figure 12I)**. These together suggest that function of *OsPLT1* is necessary for post-embryonic establishment of proper root architecture in rice.

**Figure 5:**
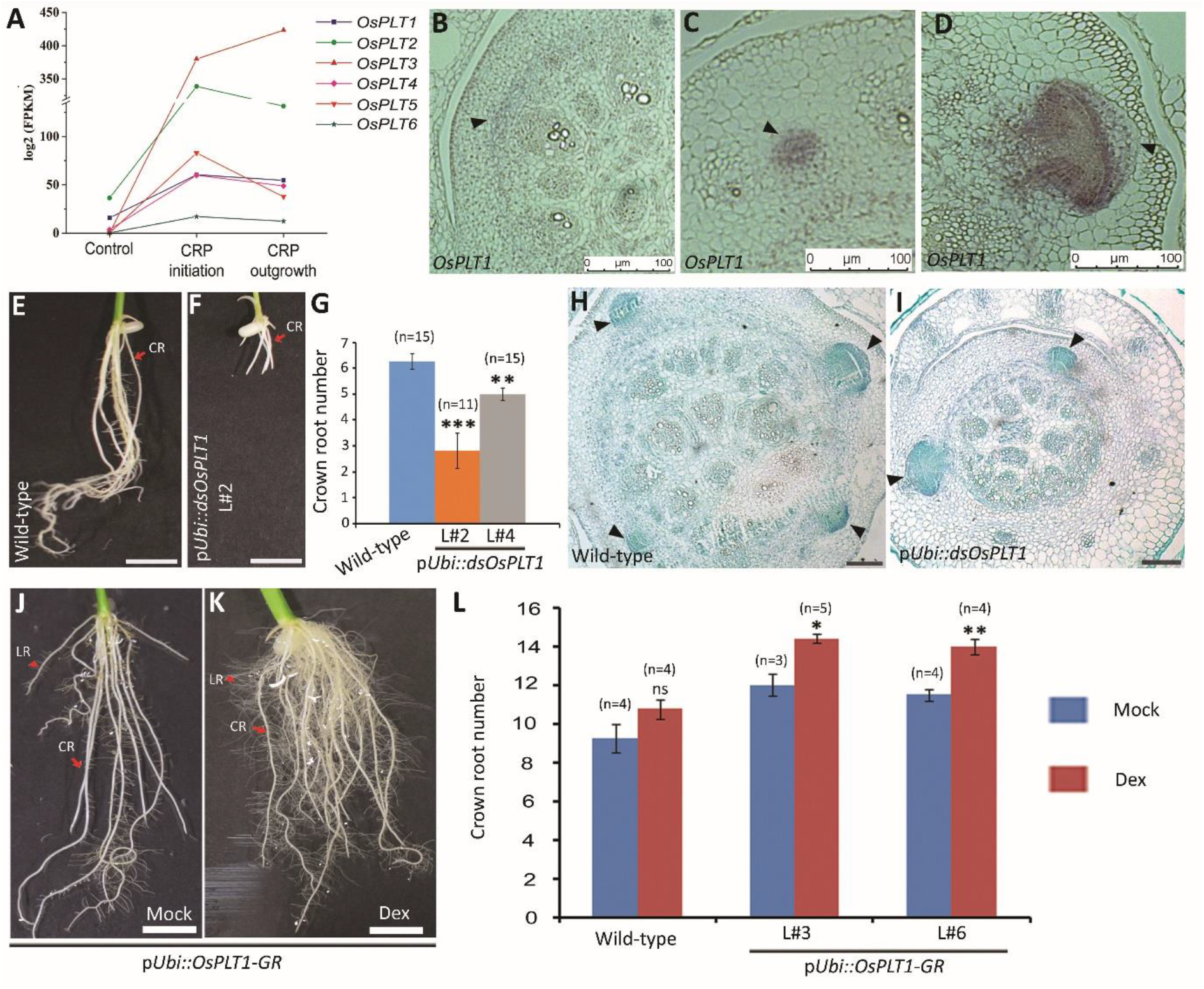
Rice *PLETHORA1* (*OsPLT1*) function is required and sufficient for post-embryonic root development. (A) LCM-seq data for the expression pattern of rice *PLETHORA* genes during CRP initiation and outgrowth. (B-D) Tissue-specific expression pattern of *OsPLT1* during rice CRP development. *OsPLT1* expression begins during CRP initiation (B, C) and continues to express in the outgrowing CRP (D). (E-G) Phenotypes of down-regulation of *OsPLT1* in p*Ubi-dsOsPLT1* rice lines. Overall root architecture is altered in knock-down lines (F), as compared to control plants (E). Length of PR, CR and LR is reduced in down-regulated lines (F). (G) Quantitative representation of CR numbers in two independent down-regulated lines, as compared to wild-type plants (**p≤0.005; ***p≤0.001; two-sample t-test). (H, I) Cross section of rice coleoptile base showed lesser CRP in the *OsPLT1* knock-down line (I), as compared to wild-type (H). (J, K) Induced over-expression of glucocorticoid receptor (GR) fused *OsPLT1* (p*Ubi::OsPLT1-GR*) causes increased number of CR and LR in rice (K), as compared to mock treated plants (J). (L) Quantitative representation of CR numbers in two independent p*Ubi::OsPLT1-GR* lines upon dexamethasone treatment (ns, not significant p>0.05; *p≤0.05; **p≤0.005; two-sample t-test). Site of CRP and developing CRP are highlighted with black arrowhead in (B)-(D), (H) and (I). Red arrows and arrow heads mark CR and LR, respectively in (E), (F), (J), and (K). (CRP, crown root primordia; CR, crown root; LR, lateral root). Bars= 100 µm in (B)-(D), (H), and (I); 1 cm in (E), (F), (J), and (K).

To further confirm function of *OsPLT1* for root development, we generated transgenic rice lines over-expressing *OsPLT1* translationally fused with c-terminal domain of glucocorticoid receptor (GR). This fusion allowed us to over-express *OsPLT1* in an inducible manner (**Supplemental Figure 13A-13C**). As opposed to the phenotypes observed upon *OsPLT1* down-regulation, growing *OsPLT1-GR* plants in presence of inducer, dexamethasone resulted in a more robust root architecture **(Figure 5K; Supplemental Figure 13C, 14D**), as compared to mock-treated plants (**Figure 5J; Supplemental Figure 13C, 14C)**. However, no such effect was observed in dexamethasone treated wild-type plants (**Supplemental Figure 13B; 14A, 14B**). Both PR and CRs developed high density and larger LRs upon dexamethasone induction (**Figure 5K**). Younger CRs (marked with red arrowheads) have not extensively developed LRs in mock-treated plants (**Figure 5J**) whereas these CRs displayed higher number of lengthy LRs upon dexamethasone treatment (**Figure 5K**). Similarly, CR root numbers were significantly increased in multiple independent *OsPLT1-GR* lines upon over-expression whereas effect of dexamethasone treatment did not give any significant change in the CR number (**Figure 5L**). Important to note that origin of CRs and LRs are diverged in rice, the CRs are originated from the stem tissues (shoot-borne) whereas LRs develop from the root tissues (root-borne). This observation suggests that *OsPLT1* promotes root development in rice, irrespective of their origin, thus has a conserved role during post-embryonic root development.

### *OsPLT1* Directly Activates Local Auxin Biosynthesis during Rice CR development

*PLETHORA* genes regulate local auxin biosynthesis to control phyllotaxis and vascular regeneration, by regulating expression of *YUCCA* genes in *Arabidopsis* (Pinon et al., 2013; Radhakrishnan et al., 2020). In rice, YUCCA-Auxin*-*OsWOX11 module regulates CR development as over-expression of rice YUCCA genes produce ectopic and massive CR formation *via* WOX11 pathway (Zhang et al., 2018). We therefore, quantified the expression levels of a few rice YUCCA genes in *OsPLT1* down-regulated and over-expression lines. *OsYUC3* is strongly expressed in the ground meristem tissues, prior to CRP establishment whereas *OsYUC1* is strongly induced during CRP initiation (**Supplemental Figure 3A**). The expression level of *OsYUC1* and *3* was reduced in the stem base of *OsPLT1* knock-down lines as compared to wild-type (**Figure 6A**), suggesting *OsPLT1* regulates their expression during CR development. Next, to study if *OsPLT1* is sufficient to induce expression of *OsYUC1* and *3* in the ectopic tissues, the expression level of these genes was analyzed in the leaf blade upon treating p*Ubi::OsPLT1-GR* line with dexamethasone to induce *OsPLT1* over-expression. The expression of both of these genes was induced upon *OsPLT1* over-expression (**Figure 6A**), further validating that *OsPLT1* promotes YUC-mediated local auxin biosynthesis. To further investigate if *OsPLT1* could directly regulate expression of these YUC genes, we treated p*Ubi::OsPLT1-GR* line with dexamethasone in presence of protein synthesis inhibitor, cycloheximide. Dexamethasone treatment induced expression of *OsYUC1* and *3* in presence of cycloheximide (**Figure 6A; Supplemental Figure 14E**), indicating that *OsPLT1* might be directly activating YUC genes during CRP establishment. To further confirm that *OsPLT1-*mediated auxin biosynthesis is required for CR formation, we selected a strong RNAi line where root was not developed upon *OsPLT1* down-regulation (**Figure 6B**), and supplied auxin exogenously. Auxin treatment induced rooting in these lines (**Figure 6D**), similar to wild-type plants (**Figure 6C**). These observations together confirm that auxin is the limiting factor in *OsPLT1* loss-of-function lines and a feed-back regulatory loop between auxin and *OsPLT1* is required for regulating root architecture in rice.

**Figure 6:**
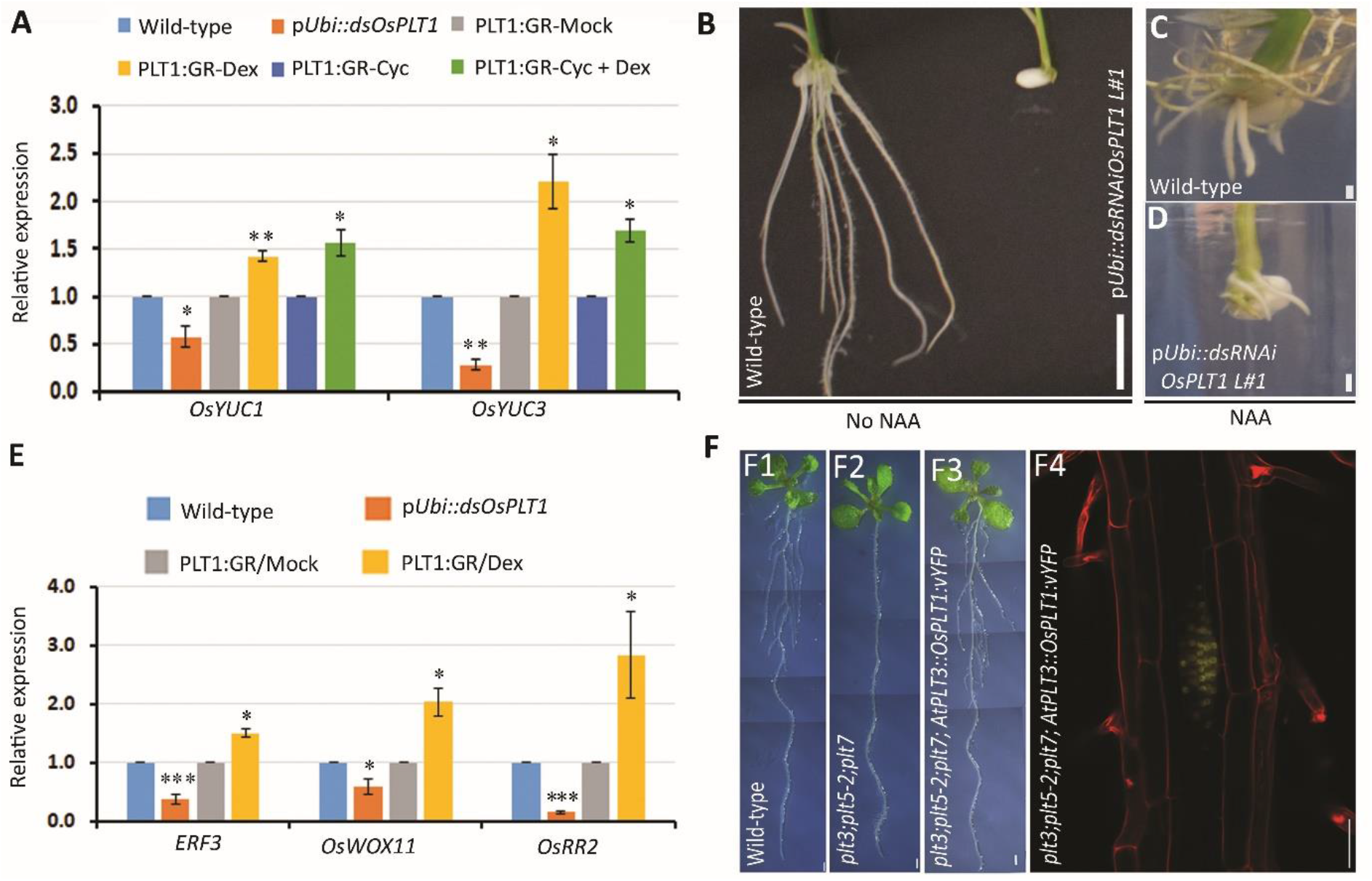
Regulatory relationship of *OsPLT1* with auxin biosynthesis and *ERF3–OsWOX11– OsRR2* pathways. (A) qRT-PCR analysis of auxin biosynthesis genes, *OsYUC1* and *OsYUC3* in *OsPLT1* down-regulated and over-expression lines. Inducible over-expression of *OsPLT1* was performed by treating leaf blades of p*Ubi::OsPLT1-GR* lines with dexamethasone, in presence (cyc + dex) and absence (dex) of protein synthesis inhibitor, cycloheximide. Ethanol (mock) and cycloheximide (cyc) were used as background controls (*p≤0.05; **p≤0.005; two-sample t-test). (B) Strong down-regulation of *OsPLT1* results in nearly complete inhibition of roots in some lines. (C, D) Exogenous auxin treatment can induce CR formation in these strong *OsPLT1* loss-of-function lines (D), similar to in control wild-type plants (C). (E) Expression levels of *ERF3, OsWOX11*, and *OsRR2* in the coleoptile base of *OsPLT1* down-regulated and dexamethose-treated p*Ubi::OsPLT1-GR* over-expression lines (*p≤0.05; ***p≤0.001; two-sample t-test). (F) Stereo images of 8-dpg wild-type plant (F1), *plt3;plt5-2;plt7* defective in LRP outgrowth (F2), and *plt3;plt5-2;plt7;AtPLT3::OsPLT1:vYFP* (F3). The rescue of LR formation in *plt3;plt5-2;plt7* mutant reconstituted with *AtPLT3::OsPLT1:vYFP* (*plt3;plt5-2;plt7;AtPLT3::OsPLT1:vYFP*). Confocal image showing expression of *OsPLT1:vYFP* in the LRP of *plt3;plt5-2;plt7;AtPLT3::OsPLT1:vYFP* (F4). Red colour in (C4) represents propidium iodide staining. (LRP, lateral root primordia; CRP, crown root primordia; CR, crown root; LR, lateral root). Bars= 1 cm in (B); 1 mm in (C), (D), (F1-F3) and 50µm in (F4).

### *OsPLT1* is an upstream regulator of ERF3-WOX11-RR2 regulatory module

A regulatory module involving ERF3, OsWOX11 and OsRR2 plays a crucial role during crown root development (Zhao et al. 2009, 2015). ERF3 physically interact with OsWOX11 and repress the expression of *OsRR2* during crown root formation (Zhao et al. 2015). We, therefore, analyzed expression levels of *ERF3, OsWOX11* and *OsRR2* in down-regulated and over-expression lines of *OsPLT1* by qRT-PCR analysis. We observed that expression of all these genes was reduced in the RNAi lines as compared with wild-type (**Figure 6E**). This suggests that *OsPLT1* regulates expression of *ERF3, OsWOX11* and *OsRR2* in the stem base.

Next, to further confirm regulation of ERF3-OsWOX11-OsRR2 module by *OsPLT1*, we analyzed expression of these genes in the stem bases of the p*Ubi::OsPLT1-GR* lines upon over-expression of *OsPLT1*. We observed opposite expression pattern of these genes upon induced over-expression of *OsPLT1* (**Figure 6E)**. The expression of *ERF3, OsWOX11* and *OsRR2* was induced upon dexamethasone treatment as compared to mock-treated plants (**Figure 6E**). These observations together suggest that *OsPLT1* functions *via* ERF3-OsWOX11-OsRR2 signaling pathway and is an upstream regulator of this regulatory module.

### Root Outgrowth Promoting Function of *OsPLTs* is Conserved in *Arabidopsis*

In *Arabidopsis, PLTs* regulate lateral root outgrowth (Du and Scheres, 2017). We next asked if function of *OsPLT1* is conserved in *Arabidopsis*. Towards this we delivered the regulator of shoot-borne crown root primordia (CRP), the rice *OsPLT1*, in the transcription domain of *Arabidopsis* lateral root primordia (LRP) of *plt3;plt5-2;plt7* triple mutant which were defective LRP outgrowth (**Figure 6F2**). Strikingly, the lateral root outgrowth defect in the *plt3;plt5-2;plt7* triple mutant was rescued to wild-type level by *OsPLT1* when it was expressed in *Arabidopsis PLT3* domain (**Figure 6F-6F4**). This suggest that *PLT-*like genes have acquired species-specific expression domain while the function of proteins is conserved i.e., to promote the outgrowth of root primordia irrespective of their developmental origin. Thus to summarize, we propose a regulatory model suggesting that auxin regulated expression and CRP-specific activation of *OsWOX10* and *OsPLT1* is required for post-embryonic adventitious and lateral root development through conserved regulators (**Figure 7**). This study has also provided an indication that expression of *OsWOX10* and *OsPLT1* might also be regulated epigenetically. Further epigenomics based studies will be required to reveal upstream regulation of *OsWOX10* and *OsPLT1*.

**Figure 7:**
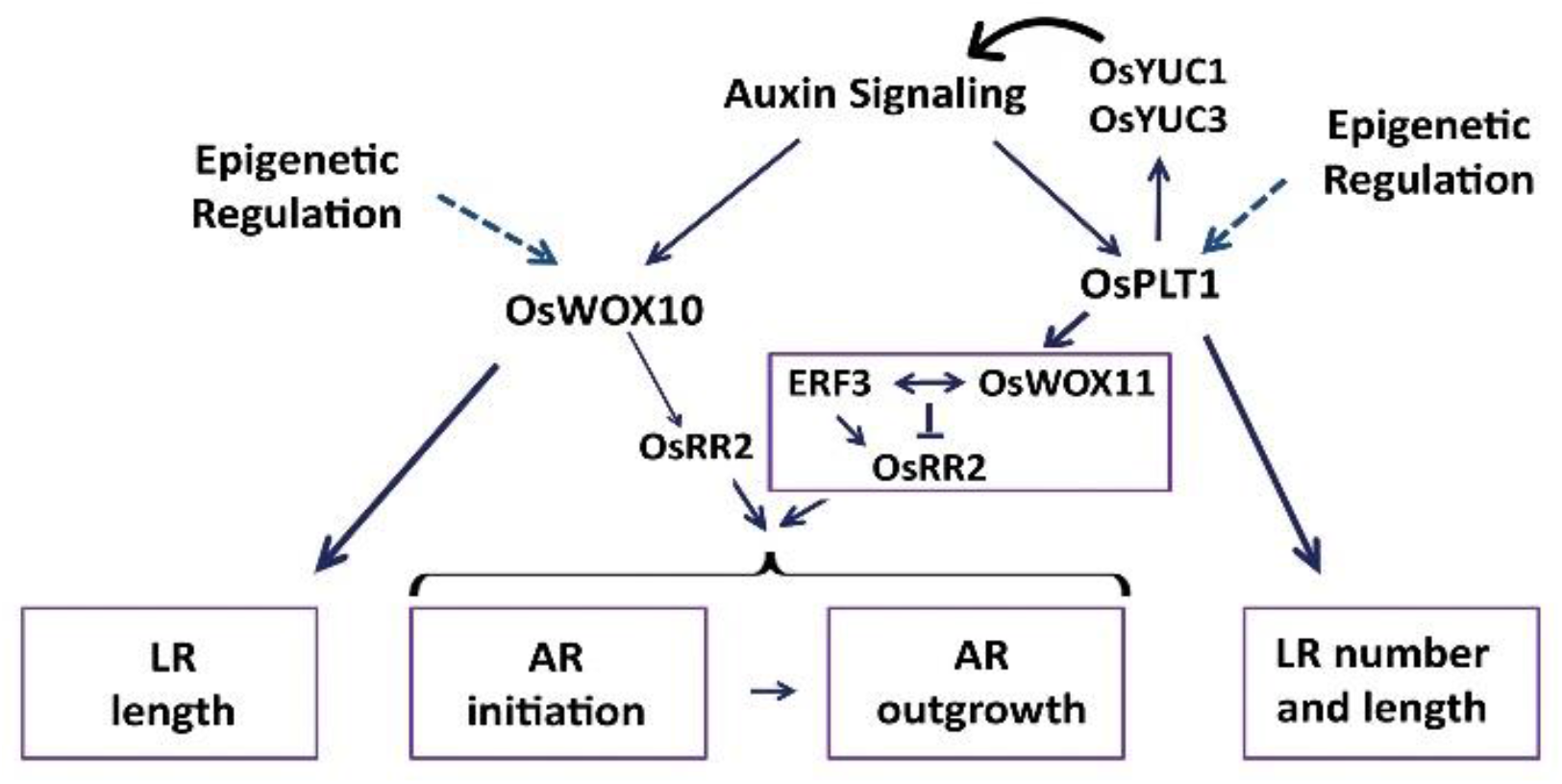
Schematic model depicting that auxin inducible expression of *OsWOX10* and *OsPLT1* in the incipient CRP promotes initiation and outgrowth CRP, and also growth of emerged CRs. *OsPLT1*, in turn directly regulate auxin biosynthesis YUC genes, thus generating a feed-back regulatory loop. *OsPLT1* also functions as an upstream regulator of ERF3-WOX11-RR2 regulatory module. Apart from regulating CR development, *OsPLT1* also promotes LR formation.

## DISCUSSION

The process of CRP development begins with an induction phase where innermost ground tissues of rice stem base re-enter the cell cycle in a localized domain to establish founder cells for CRP (Itoh et al., 2005; Guan et al., 2015). This would require a genetic reprogramming of the cells competent for CRP formation. Our study provides a global gene expression map of the CRP initiation and outgrowth and reveals the regulatory role of key root fate determinants during CRP development. We reveal how transcriptional state of genetic and epigenetic regulators in heterogeneous cellular environment coordinate the initiation and the outgrowth of CRP. Furthermore, our findings lead to discovery of previously unrecognised role of rice WOX and PLT genes in controlling the root architecture.

### Epigenetic Regulation During CRP Development

The induction phase during CRP establishment would require the acquisition of pluripotency through genetic reprogramming in response to localized developmental cues. Stable but reversible modification of DNA and histone proteins along with chromatin remodeling factors provide epigenetic regulation of gene expression to control the crucial balance between stem cell self-renewal, cellular patterning, and tissue-specific differentiation during plant growth and development, and various stresses (Takatsuka and Umeda, 2015; Servet et al. 2010; Ojolo et al. 2018; Singh et al., 2020). *Arabidopsis* GCN5-related N-acetyltransferases family (GNAT), SWI2/SNF2 factors, and PHD domain-containing factors play a key role in positioning the SCN by epigenetically regulating expression domain of *PLTs* and *WOX5* (Kornet and Scheres, 2009; Servet et al., 2010; Sang et al., 2012; Napsucialy-Mendivil et al., 2014; Zhang et al., 2015). In rice, WOX11 recruits the GCN5-ADA2 complex to activate the expression of genes required for cell division in the CR meristem (Zhou et al., 2017). A global epigenetic control of auxin signaling is evident from differential H3K27me3 pattern associated with genes involved in auxin biosynthesis, distribution, and signaling between dividing and differentiated cells or during the acquisition of pluripotency (Lafos et al., 2011; He et al., 2012; Chen et al., 2016; Yamamuro et al., 2016; Mateo-Bonmatí et al., 2019). Members of CHD3 family, *PICKLE* (*PKL*) regulates LR development through the IAA14-ARF7/19 auxin signaling module (Fukaki et al., 2006) and *OsCRL6/OsCHR4* controls CR development by epigenetically regulating expression of YUCCA genes (Wang et al., 2016; Guo et al., 2019). Our genome-wide LCM-seq analysis together with early activation of potential epigenetic modifiers in the CRP founder cells and defects in CR development in response to the perturbations in epigenetic modifications suggest its important role in trans-differentiation.

### Auxin-Transcription Factors Cross-talk Generates Gene Regulatory Modules During CRP Development

Consistent with known role of auxin maxima and auxin signaling in *Arabidopsis* post-embryonic root development, we find localized auxin maxima, progressive surge of local auxin biosynthesis genes and converse pattern of negative regulators of auxin signaling during CRP development. Different regulatory modules involving components of the auxin signaling pathway and transcription factors are known to regulate post-embryonic root development in plants. The IAA28–ARF5, 6, 7, 8, 19-GATA23 module is important for the specification of the LR founder cells whereas IAA14-ARF7, 19-LBD16, 18, PUCHI module is required for regulating LR initiation in *Arabidopsis* (De Rybel et al., 2010, Lavenus et al., 2013, Kang et al., 2013). In growing roots, the IAA17/AXR3-ARF10/16 module functions in the root meristem to restrict expression of *WOX5* to QC and to regulate the expression of *PLT1* (Ding and Friml, 2010). We observed dynamic expression pattern of related genes during rice CRP development with plausible regulatory divergence. Based on the reported functions of the putative homologous genes in LR development, their phylogenetic relationship and co-expression pattern in our LCM-seq analysis, we are tempted to speculate that OsIAA13/30-OsARF16/19-OsWOX10, OsCRL1 could be a possible regulatory module functional during CRP initiation and OsIAA11/23/30-OsARF8/22-OsDH1 could function during CRP outgrowth (Jain et al., 2006; Wang et al., 2007; Jun et al., 2011, Kitomi et al., 2012). However, OsIAA30-OsARF10-OsCRL1, QHB, OsPLTs might functions during meristem maintenance (Lavarenne et al., 2019). A detailed protein-protein interaction and combinatorial mutant analysis should demonstrate species and root-type specificity of these regulatory modules.

### *OsWOX10* Regulates the Development of Adventitious Roots

Auxin-responsive *Arabidopsis WOX11* and *WOX12* redundantly regulate first-step cell fate transition from competent cell to root founder cells during *de novo* root organogenesis (Liu et al., 2014). Our functional studies along with auxin induced activation of *OsWOX10* in the CRP founder cells uncovers the non-redundant function of *OsWOX10* in priming developmental program for root organogenesis during plantlet regeneration and vegetative development. Among other related WOX members, *OsWOX11* is known to regulate CR initiation and growth, as loss-of *wox11* function fails to develop CRs and its over-expression causes stimulated and ectopic root production (Zhao et al., 2009). *OsWOX12* (*OsWOX6*) is expressed in the developing CRP (Neogy et al., 2019) and has been shown to be involved together with *OsWOX11* in regulating tiller angles in rice (Zhang et al., 2018). We cannot rule out the possibility of genetic redundancy among these members in regulating AR development. A combinatorial mutant analysis will be required to study genetic interactions among these factors.

Though ARs are major component of root architecture in monocot plants, dicot species also develop ARs, mainly in response to specific environmental signals and wounding but their origin is diversified (Bellini et al., 2014). In *Arabidopsis*, the pericycle cells at the xylem pole of hypocotyl give rise to ARs. We showed that the function *OsWOX10* is conserved in promoting ARs from hypocotyl of *Arabidopsis* plants. This suggests that *OsWOX10* can activate correct genetic program of ARs in the competent tissues, irrespective of the plant species.

### Function of *OsPLT1* is Conserved in Controlling Post-embryonic Root Development

Root architecture and origin of post-embryonic roots are diverged across the plant species. In *Arabidopsis*, both ARs and LRs originate from the xylem pole pericycle cells of hypocotyl and primary root, respectively whereas, in rice, ARs/CRs are developed from the innermost ground tissues of shoot (Itoh et al. 2005, Rebouillat et al. 2009, Bellini et al. 2014). Thus, the developmental context of LR in *Arabidopsis* and CR in rice is distinct. *OsPLT1* regulates initiation and growth of all post-embryonic roots, as the number and length of CRs and LRs were inversely affected in knock-down and over-expression rice lines. Functional conservation of rice root fate determinant is further evident from rescue of root pericycle originated LRPs outgrowth in *Arabidopsis plt3,5,7* mutant by delivering *OsPLT1* in LRP transcriptional domain. It is likely that conserved root promoting factors such as PLTs have acquired species specific function in the two evolutionary diverged plants species, rice and *Arabidopsis*, largely by modulating the cis-regulatory sequences but not the function of protein. Taken together, our studies provide CRP stage-specific temporal gene expression map, and reveal the function of OsPLT1-dependent transcription regulatory module in controlling the rice root architecture.

Recent years have seen the burst of literature demonstrating the role of WOX and PLT genes in controlling the organ primordia initiation and their outgrowth both, during embryonic and well as post-embryonic development of *Arabidopsis* (Costanzo et al., 2014, Horstman et al., 2014, Dolzblasz et al., 2016; Bustillo-Avendaño et al., 2018). Here, we show their role in controlling shoot-borne root primordia development in rice. It is tempting to speculate that primary role of these genes in controlling the organ primordia development is conserved across the plant species. It will be interesting to see how these conserved transcription factors have acquired organ specific function in different plant species.

### A Potential Regulatory Feedback Loop Controls CRP Development

Local auxin biosynthesis and polar auxin transports generate auxin maxima at the site of root primordia, prior to their establishment (Benkova et al., 2003; De Rybel et al., 2010). The expression of *OsPLT1* is induced by auxin treatment (Li and Xue, 2011). Interestingly, our study shows that *OsPLT1* can activate local auxin biosynthesis by upregulating the expression of *OsYUC* genes. This potential auxin-*OsPLT1* positive regulatory feedback loop appears essential to establish the competence in shoot cells and initiate CRP-specific developmental program (**Figure 7**).

In rice, a regulatory interaction of *ERF3, OsWOX11* and *OsRR2* plays a crucial role during initiation and emergence of CRs (Zhao et al., 2015). Recently, YUCCA-auxin-OsWOX11 module has been shown to function during CR development (Zhang et al., 2018). Furthermore, we show that *OsPLT1* functions upstream of *ERF3-OsWOX11-OsRR2* signaling pathway. Taken together, our studies provide the possibility of a tangled regulatory loops between auxin and conserved regulators of root development (**Figure 7**). It is tempting to speculate that multiple regulatory feedback loops between root specific transcription factors and plant hormones shape up the architecture of shoot borne root in evolutionary diverged grass species.

## METHODS

### Plant Material, and Treatments

*Oryza sativa* seed germination, growth, auxin treatment, sample collection and rice transformation were performed as described by Neogy et al., (2019). Sodium butyrate (HiMedia, India), and sodium acetate (HiMedia, India) were dissolved in water whereas 5-azacytidine (HiMedia, India) was dissolved in dimethyl-sulfoxide (DMSO). The surface sterilized, rice IR-64 seeds were grown on ½ MS media with or without 150 µM sodium butyrate, 3 mM sodium acetate, and 20 µM 5-azacytidine. For inducible down-regulation of *OsWOX10*, 10 µM 17β-estradiol (Sigma-Aldrich), dissolved in DMSO, was used. For inducible over-expression of *OsPLT1*, 10 µM dexamethasone (Sigma-Aldrich), and/or 10 µM cycloheximide (Sigma-Aldrich), both dissolved in ethanol, were used. Cycloheximide treatments were started 30 min before the dexamethasone treatment in the samples that were treated with both reagents. For auxin treatment in strong *dsplt1* RNAi lines, seedlings were grown on ½ MS media for 9 days along with WT control. All old roots were cut and new roots were induced on ½ MS media supplemented with 0.1mg/L NAA. The *Arabidopsis* mutants *wox11-2* (SALK_004777), *wox12-1* (SALK_087882) and *wox11-2 wox12-1* seeds were kindly provided by Dr. Lin Xu used by Liu et al., (2014).

### Laser Capture Microdissection

For LCM, the 1mm coleoptile base tissue from 6-day old rice seedling (var. IR-64) was harvested in Carnoy’s fluid (ethanol: chloroform: acetic acid glacial; 6:3:1), infiltrated twice under mild vacuum and dehydrated through graded ethanol series followed by replacement with xylene. The tissue was embedded in Paraplast (Sigma-Aldrich) and 8-µm thin sections were cut using RM2125 microtome (Leica) and sections were taken on PEN membrane slides (Carl-Zeiss, Germany). The CRP were micro-dissected on PALM Microbeam (Carl-Zeiss) and collected in RNA extraction buffer.

### RNA Extraction, Library Preparation and RNA Sequencing

RNA was isolated from LCM collected CRP using ARCTURUS PicoPure RNA Isolation Kit (ThermoFisher, Waltham, MA, USA) according to manufacturer’s protocol. The extracted RNA was assayed for RNA integrity on 2100 Bioanalyzer using RNA 6000 Pico Kit (Agilent Technologies, Santa Clara, CA, USA). RNA samples were depleted for ribosomal RNAs (rRNAs) using Ribo-Zero rRNA Removal Kit (Illumina, San Diego, CA, USA) and were used for cDNA synthesis and library preparation using SMARTer universal low input RNA kit (Clontech, Mountain View, CA). Library fragment size distribution was checked on the Agilent 2100 Tapestation System with the High Sensitivity D1000 Kit (Agilent Technologies). A total of six RNA sequencing libraries were sequenced on Nova Seq 6000 Platform (Illumina).

### Sequence Alignment and Gene Cluster Analysis

The sequenced paired-end reads were mapped to a reference genome (MSU release 7) using STAR (Dobin et al. 2013) in two-pass mode. A gene count matrix was generated using the quant mode-GeneCounts with rows corresponding to individual genes and columns corresponding to samples. For common expression pattern analysis, fuzzy c-means clustering was performed on the data using Mfuzz (Kumar and Futschik, 2007). The gene count table was made homoscedastic using the variance stabilizing transformation function from DESeq2 (Love et al., 2014). The biological replicates for each stage were collapsed using the geometric mean of the two values. After running fuzzy c-means, a membership cutoff of 0.5 was used for assigning genes to individual clusters. The number of clusters was empirically determined by studying the Principal Component Analysis (PCA) plots, minimum cluster centroid distance, and normalized expression plots, with the number of clusters varying from 2 to 25 as in (Harrop et al., 2016). Cluster-wise gene list (log2 fold enrichment ≥ 1; p< 0.05) was used to perform GO enrichment analysis using monocot PLAZA 4.5 workbench (Bel et. al., 2018).

### Differential Gene Expression and Gene Ontology Analysis

The count matrix was used as input for differential expression analysis using DESeq2. Genes with an adjusted p-value (or q-value) less than 0.05 and the log2 fold-change ≥ 1 or ≤ −1 were considered differentially expressed genes (DEGs). For auxin responsiveness, genes with p-value less than 0.05 and the log2 fold-change ≥ 0.9 were considered as auxin inducible. Gene expression in different root zones was analyzed using CoNekT database (Proost and Mutwil, 2018; https://conekt.sbs.ntu.edu.sg/heatmap/) and heatmap was generated using tool Heatmapper (Babicki et al. 2016; http://heatmapper.ca/expression/). GO enrichment analysis was performed using BiNGO plug-in of Cytoscape (version 3.3.0) with P-value ≤0.05. GO enrichment of each condition was further used to make a comparative enrichment map via Cytoscape.

### RNA Extraction and Real-Time PCR

Total RNAs was extracted from crown tissues, and seedlings using RNeasy Plant Mini Kit (Qiagen, Hilden, Germany) followed by elimination of DNA using on-column DNase (Qiagen) according to manufacturer’s protocol. The cDNA synthesis and qRT-PCR was performed as described earlier (Neogy et al., 2019) using iScript cDNA synthesis kit and iTaq Universal SYBR Green Supermix (Bio-Rad Laboratories, India). Rice *UBQ5-*normalized ΔΔCt was used to calculate log2 fold change. A list of primers is provided as Supplemental Table 3. For statistical significance of fold-change, p-values were calculated using student t-test for two samples assuming unequal variances from three replicates.

### RNA-RNA *in situ* Hybridization

For preparing anti-sense DIG-UTP-labeled riboprobes, 150 bp of *OsERF3*, 186 bp of *OsARF16*, 170 bp of *OsHOX1*, 121 bp of *OsTrx1*, and 132 bp of *OsSWIB/MDM2* gene-specific fragments were cloned in *p*Bluescript SK+ (anti-sense), linearized with *EcoRI* and transcribed with T7 RNA Polymerase (Sigma-Aldrich). Their sense clones, linearized with *HindIII* was used to generate sense probes. The 193 bp of *OsRR24* and 134 bp of *OsATXR6* gene-specific fragment was cloned in pBS SK+ (anti-sense), *EcoRI/*T7 RNA polymerase and *HindIII/*T3 RNA polymerase (NEB) generated antisense and sense probes, respectively. For YFP anti-sense probe, 609 bp of YFP fragment cloned in pBS SK+ (anti-sense), was transcribed with *EcoRI/*T7 RNA polymerase. For *OsWOX10*, 69 bp fragment cloned in pBS SK+ (sense), *HindIII/*T3 RNA polymerase was used to generate anti-sense probes. For *OsPLT1* anti-sense probe, 617 bp of the gene specific region was cloned in pBS SK+ (sense) and linearized using *NdeI*. Both YFP and *OsPLT1* anti-sense probe were hydrolyzed to about 100 bp before use. Hybridization and detection was performed on cross-sections, as described by Neogy et al., (2019; 2020).

### Immunohistochemistry

For immunohistochemistry, tissues were treated for antigen retrieval in antigen retrieval buffer (10mM Tris, 1mM EDTA, pH-9.0). Slides were blocked with 1% BSA in 1XTBST and incubated with anti-GFP primary antibody in 1:500 dilutions (Rockland Immunochemicals, cat. no. 600-301-215S-Anti GFP mouse monoclonal antibody) for 10-12 hours. Slides were washed with 1xTBST and incubated with HRP tagged secondary antibody in 1:3000 dilutions. Colour detection was done using 3, 3′-Diaminobenzidine (Sigma-Aldrich) as a substrate. Sections were counterstained with Hematoxylin, dehydrated with graded ethanol, cleared with xylene and mounted in DPX.

### Plasmid Construction and Generating Transgenic Line

For generating DR5ev_erYFP transgenic rice lines, the DR5ev_erYFP-nosT (1.4kb) was PCR amplified using the plasmid pHm-DR5ev-erYFP_nosT2 as template (gift from Ari Pekka Mahonen’s lab at Helsinki University, Finland) and cloned into pCAMBIA1390 vector. For *OsWOX10* down-regulation construct, a 526 bp gene-specific fragment was used to generate inverted repeat RNAi hairpin loop and was cloned in a vector expressing estradiol-inducible XVE under maize Ubiquitin promoter. For ectopic over-expression, full length CDS of *OsWOX10* was cloned under maize Ubiquitin promoter. For *OsPLT* RNAi construct, gene-specific fragment of *OsPLT1* (979 bp) was used and dsRNAi*OsPLT1* was expressed under maize Ubiquitin promoter. For generating OsPLT1-GR construct, full-length CDS of *OsPLT1* without stop codon (1.47 kb) was PCR amplified and was cloned in pUGN vector for translational fusion with the rat glucocorticoid receptor (Prasad et al., 2005). These constructs were mobilized to *Agrobacterium tumefaciens* LBA4404 and used to raise transgenic rice lines as described by Toki et al. (2006) and Neogy et al. (2019). For *OsWOX10* over-expression in *Arabidopsis*, full-length cDNA was PCR amplified and cloned under *Ubiquitin 10* promoter. These constructs were introduced into GV3101 *Agrobacterium* by electroporation and transformed into *Arabidopsis wox11-2, wox12-1, and wox11-2/wox12-1* mutant plants by floral dip method (Clough and Bent, 1998). For complementation of *Arabidopsis plt* mutants, *OsPLT1 (LOC_Os04g55970*.*2)* was amplified from genomic DNA extracted from *Oryza sativa* leaf tissues. The *OsPLT1* gene was cloned under *Arabidopsis PLT3* promoter (7.7Kb) and tagged with v*YFP*. (Radhakrishnan et al., 2020). The constructs for *Arabidopsis* transformation were cloned using Multisite gateway recombination cloning system (Invitrogen) using pCAMBIA 1300 destination vector. These constructs were introduced into C58 *Agrobacterium* by electroporation and transformed into *Arabidopsis plt3; plt5-2; plt7* mutant plants by floral dip method (Clough and Bent, 1998).

### Rice phenotyping

For studying phenotype of *OsWOX10* down-regulation on root regeneration, healthy regenerated shoot with no roots were transferred from shoot regeneration media supplemented with 50 mg/L hygromycin B to root induction media (Toki et al., 2006) containing 50 mg/L hygromycin B, DMSO (mock) or 10 µM 17ß-estradiol. The number of regenerated roots were counted on the 11, 13 and 20 days post-induction. For *OsWOX10* overexpression and *OsPLT1* constitutive downregulation, the seeds were grown as described by Neogy et al., (2019). The roots were counted and stem base collected for RNA. For studying CR phenotype of *OsWOX10* down-regulation, seeds were grown on ½ MS media supplemented with 10 µM 17ß-estradiol and DMSO as control. *OsPLT1*-GR transgenic seeds were surface sterilized and germinated on ½ MS media supplemented with 0.1% ethanol as mock or 5 µM dexamethasone. After eight days, CR number was counted in both wild-type and transgenic lines. For statistical significance of CR number, p-values were calculated using student t-test for two samples assuming unequal variances.

### Accessions Numbers

Sequence data from this article can be found in RiceXPro/TAIR database under the following accession numbers: *OsWOX10, LOC_Os08g14400*; *OsWOX11, LOC_Os07g48560*; *OsWOX12, LOC_Os03g20910*; *OsPLT1, LOC_Os04g55970*.*2*; *OsPLT2, LOC_Os06g44750*.*1*; *OsPLT3, LOC_Os02g40070*; *OsPLT4, LOC_Os04g42570*; *OsPLT5, LOC_Os01g67410*; *OsPLT6, LOC_Os11g19060*; *OsTRX1, LOC_Os09g04890*; *OsSWIB/MDM2, LOC_Os03g55310*; *OsATXR6, LOC_Os01g73460*; *OsERF3, LOC_Os01g58420*; *OsARF16, LOC_Os06g09660*; *OsHOX1, LOC_Os10g41230*; *OsRR2, LOC_Os02g35180*; *OsRR24, LOC_Os02g08500. OsYUC1, LOC_Os01g45760*; *OsYUC3, LOC_Os01g53200*; *AtPLT3, At5g10510*; *AtPLT5, At5g57390*; *AtPLT7, At5g65510*.

## Author Contributions and Acknowledgments

T.G. performed experiments of LCM, RNA *in situ* hybridization and *OsWOX10* function in rice. Z.S. performed experiments for auxin responses and loss-of-function of *OsPLT1* in rice. K.C. generated and analyzed *OsPLT1* over-expression lines. A.K.D., R.S.S and M.J. performed RNA sequencing data analysis. M.Y. with T.G. contributed for function of *OsWOX10* in *Arabidopsis*. V.V. and K.P. studied function of *OsPLT1* in *Arabidopsis*. K.K.K.M. cloned gene fragments for generating ribo-probes. D.C. contributed in LCM experiments. D.S. provided scientific inputs. M.J., K.P. and S.R.Y. designed experiments, analyzed data and wrote the manuscript. S.R.Y. acknowledges financial support from Department of Biotechnology (DBT), Government of India (grant # BT/PR13488/BPA/118/105/2015). Tata Innovation Fellowship to M.J. from the Department of Biotechnology, Government of India, is acknowledged. K.P. acknowledges grants from the DBT (grant# BT/PR12394/AGIII/103/89½014) and DST-SERB (grant# EMR/2017/002503/PS), Government of India and also IISER-TVM for infrastructure and financial support. Indian Institute of Technology, Roorkee (IIT Roorkee) is acknowledged to provide infrastructure support to S.R.Y. and fellowships to T. G., Z. S. and K.K.K.M. Fellowships to K.C. from University Grant Commission (UGC), A.K.D. from Indian Council of Medical Research (ICMR) and to V.V. from Council of Scientific and Industrial Research (CSIR) are gratefully acknowledged. We thank Dr. Deepak Sharma, IIT Roorkee and Bencos Research Solutions Pvt. Ltd., Bangalore, India for their initial help in RNA-seq data analysis. Secondary antibodies for immunohistochemistry was gifted by Dr. Prabhat Kumar Mandal, IIT Roorkee, India. Plasmid pHm-DR5ev-erYFP_nosT2 was a gift from Dr. Ari Pekka Mahonen’s lab at University of Helsinki, Finland. Dr. Lin Xu, Shanghai Institutes for Biological Sciences, Chinese Academy of Sciences, Shanghai, China is acknowledged for kindly providing *Arabidopsis wox11-2, wox12-1* and *wox11-2 wox12-1* seeds. Ashish Kumar and Sonia Choudhary are acknowledged for their support in growing plants.

## Supplementary Data

**Supplemental Figure 1:** Gene ontology analysis of DEGs. (A) GO terms associated with genes specifically de-regulated during CRP initiation and outgrowth. (B) GO analysis of DEGs when CRP progress from initiation to outgrowth stage.

**Supplemental Figure 2:** (A-C) RNA *in situ* hybridization using sense riboprobes for *OsTRX1* (A), *OsATXR6* (B), and *OsSWIB/MDM2* (C) on cross sections of wild-type stem base. (D) Expression level of *OsWOX10* and *OsPLT1* upon pharmacological interference of epigenetic regulation (*p≤0.05; **p≤0.005; ***p≤0.001; two-sample t-test). Bars= 100 µm in (A), 250 µm in (B) and 50 µm in (C).

**Supplemental Figure 3:** Dynamic expression pattern of auxin signaling genes. (A-C) Heatmap showing expression pattern of genes involve in auxin biosynthesis (YUC genes) and homeostasis (GH3 genes) (A), Aux/IAA genes (B), and auxin response factors (C), as derived from LCM-seq data. (D) RNA *in situ* hybridization using anti-sense YFP riboprobes on wild-type plant. Bars= 100 µm in (D).

**Supplemental Figure 4:** Differentially regulated transcriptional regulators in developing CRP.

(A) Venn diagram showing common and unique differentially expressed TFs during CRP initiation and outgrowth. (B, C) Expression pattern of *OsSHRs* and *OsSCRs* (B), and *QHB* (C) genes during CRP initiation and CRP outgrowth, as derived from LCM-seq data.

**Supplemental Figure 5:** Auxin responsiveness of differentially expressed transcriptional regulators in developing CRP. (A-C1) Expression pattern and auxin induction of TFs, activated exclusively during CRP initiation (A and A1), specifically during CRP outgrowth (B and B1), and in both the stages (C and C1). Heatmaps in (A), (B), and (C) are derived from LCM-seq data whereas plots in (A1), (B1), and (C1) are derived from auxin-responsive RNA sequencing data published by Neogy et al., (2019).

**Supplemental Figure 6:** Selected auxin-responsive CRP-expressed TFs for validation (A-C) Expression pattern of TFs during CRP development from LCM-seq data (D) Validation of these TFs for their auxin responsiveness by qRT-PCR analysis.

**Supplemental Figure 7:** (A-D) RNA *in situ* hybridization using sense riboprobes of *OsERF3* (A), *OsRR24* (B), *OsARF16* (C), and *OsHOX1* (D), on cross sections of stem base of wild-type plant. Bars= 100 µm in (A)-(D).

**Supplemental Figure 8:** Effects of 17-β estradiol treatment in wild-type and *OsWOX10* down-regulation. (A, B) Plant morphology of wild-type (A) and pUbi::XVE>dsRNAi*OsWOX10* lines upon mock and 17-β estradiol treatment. No significant effect was observed in wild-type plants. Bars= 1cm in (A) and (B).

**Supplemental Figure 9:** Effects of *OsWOX10* down-regulation. (A) Root numbers are significantly reduced on 20 days after root induction of regenerating pUbi::XVE>dsRNAi*OsWOX10* lines upon estradiol treatment in the rooting media. (B-F) Plant morphology and root architecture when *OsWOX10* is down-regulated upon 17-β estradiol treatment in pUbi::XVE>dsRNAi*OsWOX10* lines in T1 generation. (G) Expression level of related WOX-genes, *OsWOX11* and *OsWOX12* upon *OsWOX10* down-regulation during CR development in T1 generation (ns, not significant; p>0.05; two-sample t-test). Bars= 1cm in (B)-(E).

**Supplemental Figure 10:** Over-expression phenotypes of *OsWOX10*. (A, B) Extensive adventitious roots were seen in regenerating pUbi-OsWOX10 lines in root induction media (RIM). (C-G) Plant morphology and root architecture upon ectopic over-expression of *OsWOX10* during rice crown root development in T1 generation. (H) Crown root number is significantly increased in pUbi::*OsWOX10* line. (I, J) Over-expression of *OsWOX10* in p*Ubi::OsWOX10* line during plant regeneration (I) and CR development (J) by qRT-PCR analysis. Bars= 1cm in (A)-(G).

**Supplemental Figure 11:** Conserved function of *OsWOX10* in promoting adventitious root formation in *Arabidopsis*. (A, B) Effects of over-expression in three week old plants, non-transformed wild-type (*col*), *wox11-2, wox12-1*, and *wox11-2 wox12-1* mutants (A) and extensive root formation upon overexpression of *OsWOX10* in these backgrounds (B). Bars= 1cm in (A) and (B).

**Supplemental Figure 12:** Functional studies of *OsPLT1* during root development. (A, B) Morphology of wild-type (A) and pUbi::dsRNAi*OsPLT1* L#2 (B) plants. (C, D) Lateral root phenotype of pUbi::dsRNAi*OsPLT1* L#2. (E-H) Root architecture phenotypes pUbi::dsRNAi*OsPLT1* L#4. (I) Expression level of PLT genes, in pUbi::dsRNAi*OsPLT1* L#2 (ns, not significant; p>0.05; ***p<0.001; two-sample t-test). Bars= 1cm in (A)-(H).

**Supplemental Figure 13:** Molecular and phenotypic characterization of *OsPLT1* over-expression lines. (A) RT-PCR analysis of *OsPLT1-GR* lines showing expression of *OsPLT1-GR* fusion transcripts. (B, C) Plant morphology of wild-type (B) and pUbi::*OsPLT1-GR* L#6 (C) upon dexamethasone treatment. No significant effect was seen on the gross morphology of wild-type plants upon dexamethasone treatment. Bars= 1cm in (B) and (C).

**Supplemental Figure 14:** Root architecture phenotypes of *OsPLT1* over-expression lines. (A-D) Root architecture was altered upon dexamethasone treatment in pUbi::*OsPLT1-GR* plants (D), as compared to mock treated plants (C) but no effect was seen on the root architecture of wild-type plants upon dexamethasone treatment (B) as compared to mock treated plants (A). (E) RT-PCR analysis of *OsPLT1-GR* fusion transcripts in the plants treated with Mock, dexamethasone alone (Dex), cycloheximide alone (Cyc), and dex and cyc together in combination. Bars= 1cm in (A)-(D).

