## Supplemental Figure 1-14 for "Genome-Wide High Resolution Expression Map and Functions of Key Cell Fate Determinants Reveal the Dynamics of Crown Root Development in Rice"

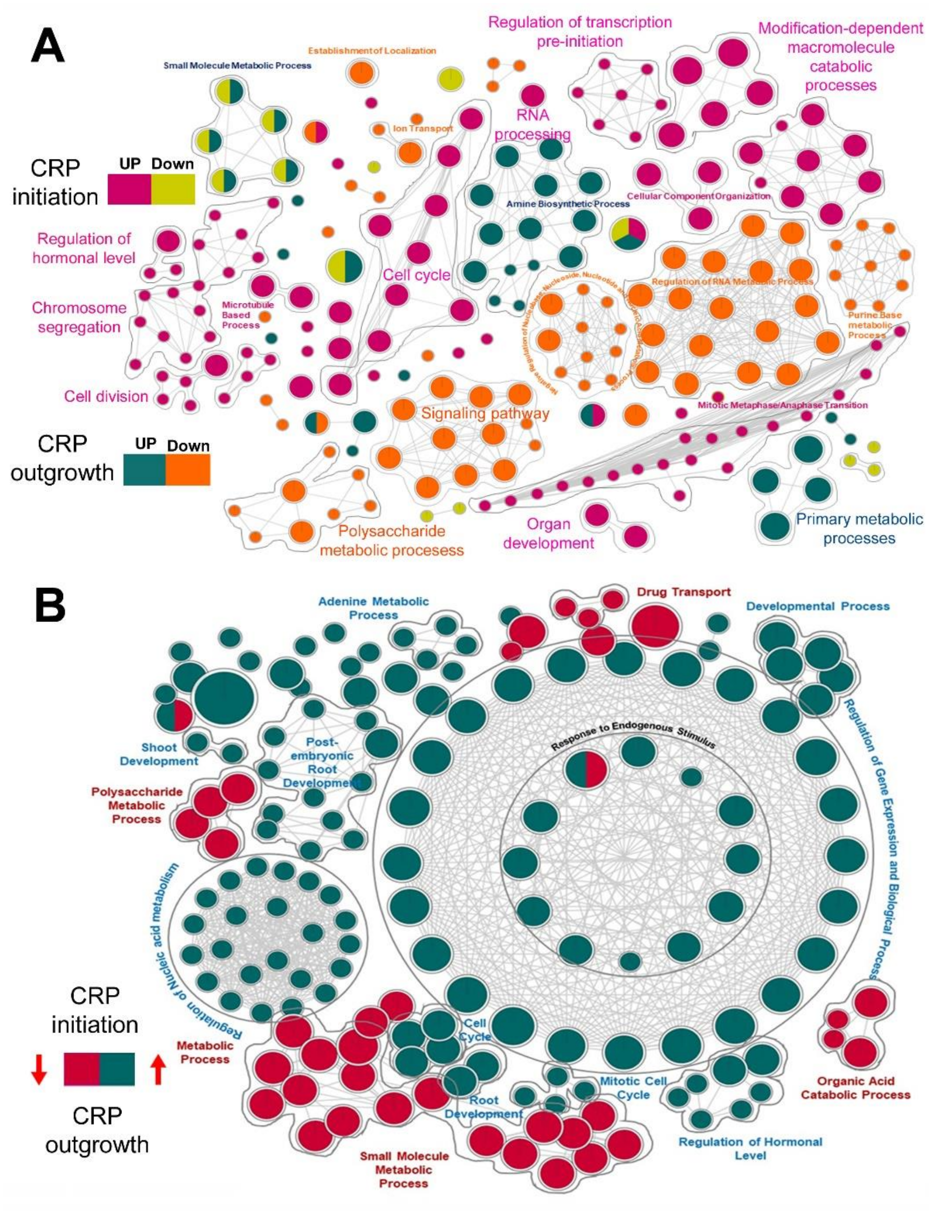


**Supplemental Figure 1:** Gene ontology analysis of DEGs. (A) GO terms associated with genes specifically de-regulated during CRP initiation and outgrowth. (B) GO analysis of DEGs when CRP progress from initiation to outgrowth stage.

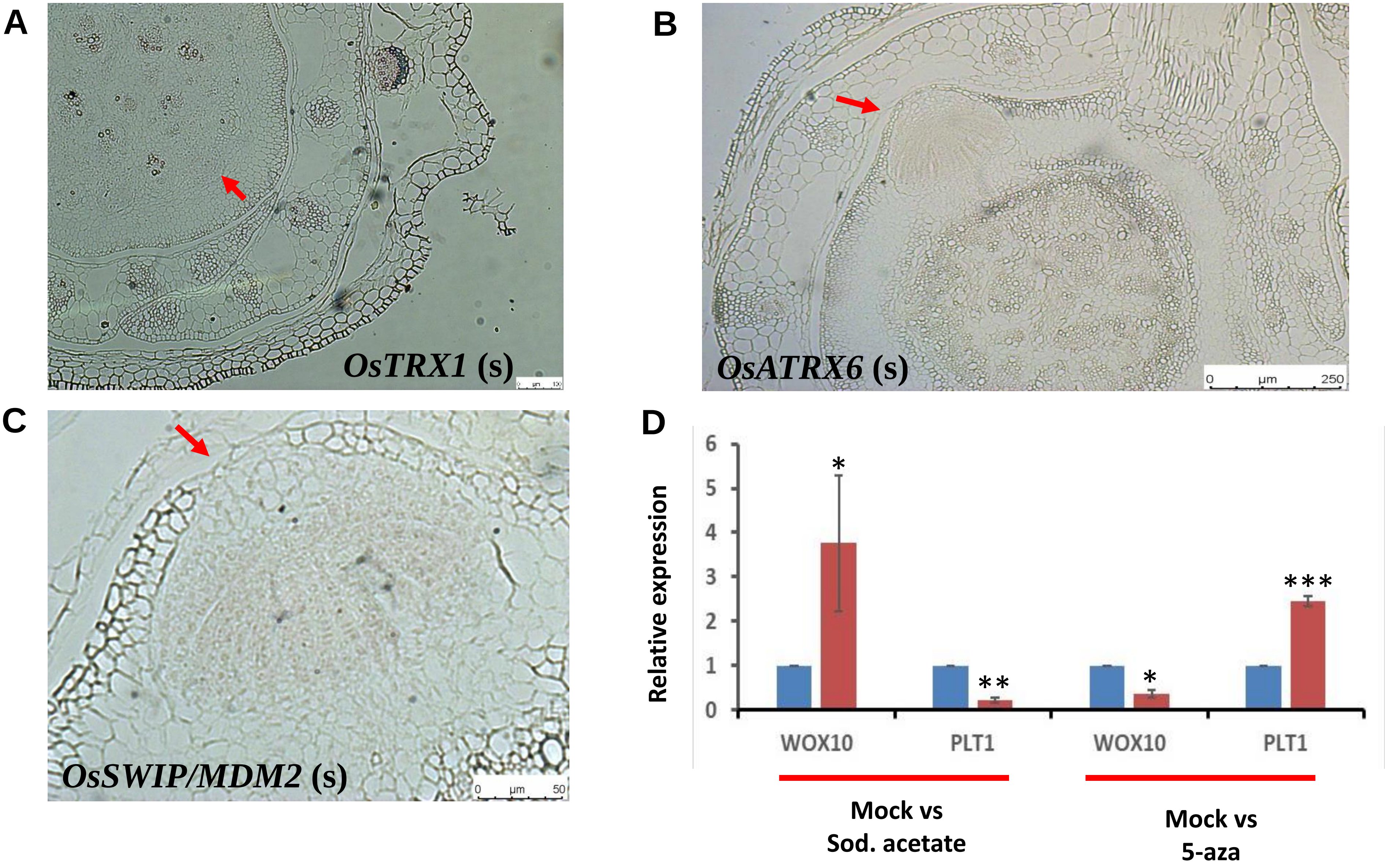


**Supplemental Figure 2:** (A-C) RNA *in situ* hybridization using sense riboprobes for *OsTRX1* (A), *OsATXR6* (B), and *OsSWIB/MDM2* (C) on cross sections of wild-type stem base. (D) Expression level of *OsWOX10* and *OsPLT1* upon pharmacological interference of epigenetic regulation (*p≤0.05; **p≤0.005; ***p≤0.001; two-sample t-test). Bars= 100 µm in (A), 250 µm in (B) and 50 µm in (C).

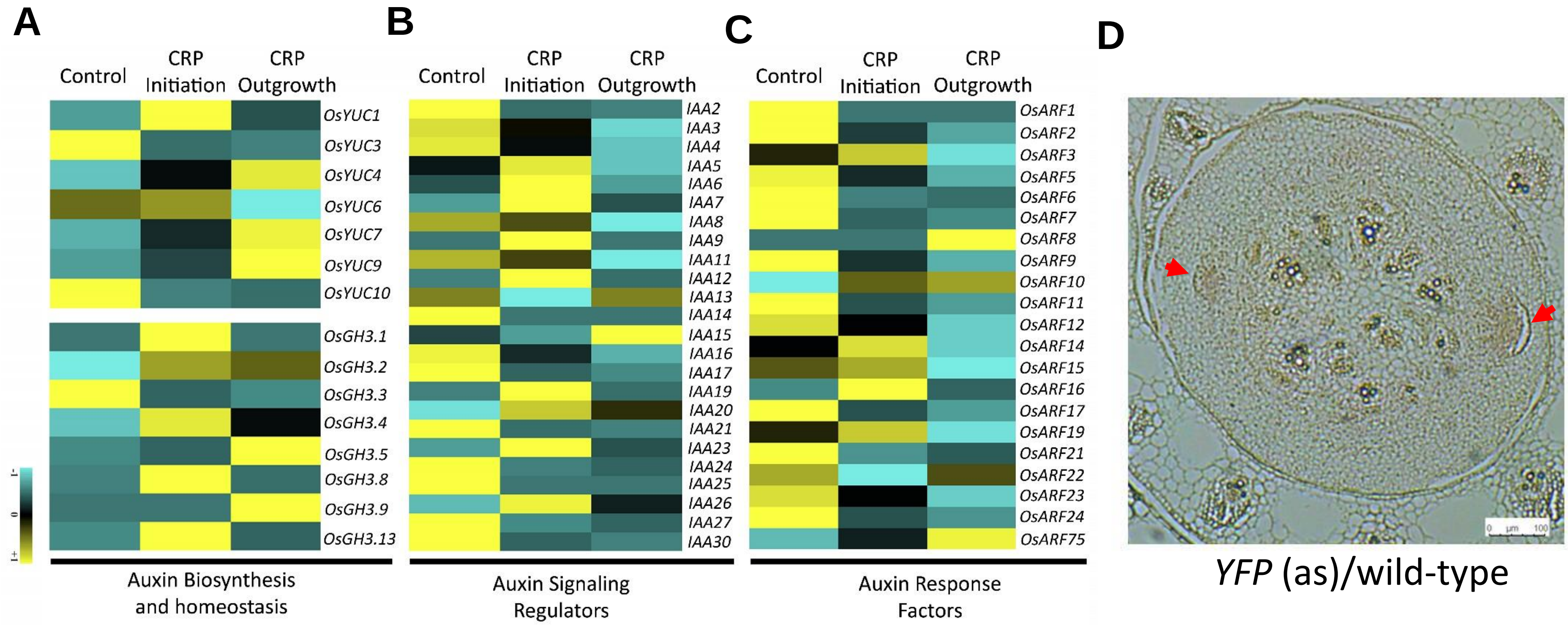


**Supplemental Figure 3:** Dynamic expression pattern of auxin signaling genes. (A-C) Heatmap showing expression pattern of genes involve in auxin biosynthesis (YUC genes) and homeostasis (GH3 genes) (A), Aux/IAA genes (B), and auxin response factors (C), as derived from LCM-seq data. (D) RNA *in situ* hybridization using anti-sense YFP riboprobes on wild-type plant. Bars= 100 µm in (D).

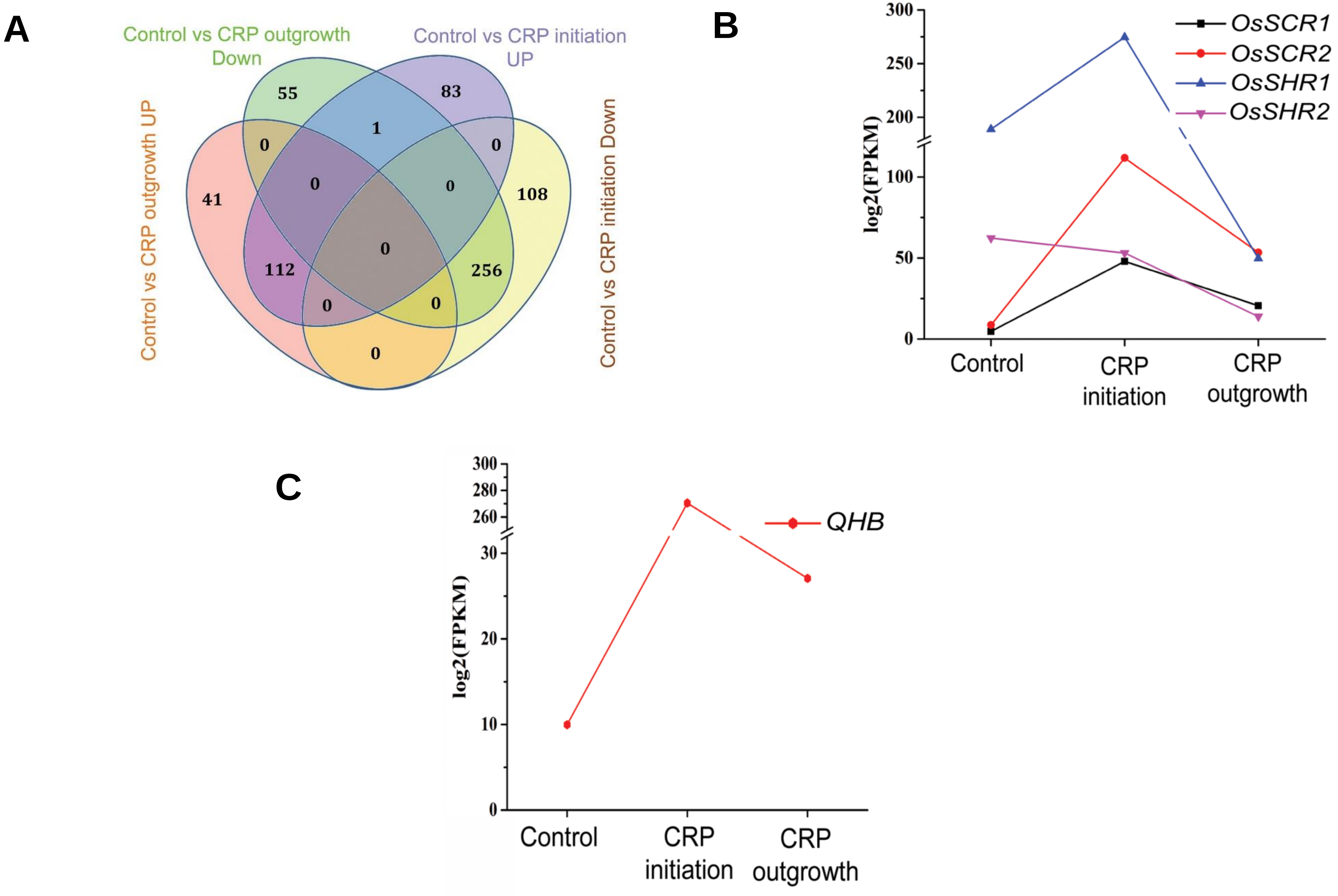


**Supplemental Figure 4:** Differentially regulated transcriptional regulators in developing CRP. (A) Venn diagram showing common and unique differentially expressed TFs during CRP initiation and outgrowth. (B, C) Expression pattern of *OsSHRs* and *OsSCRs* (B), and *QHB* (C) genes during CRP initiation and CRP outgrowth, as derived from LCM-seq data.

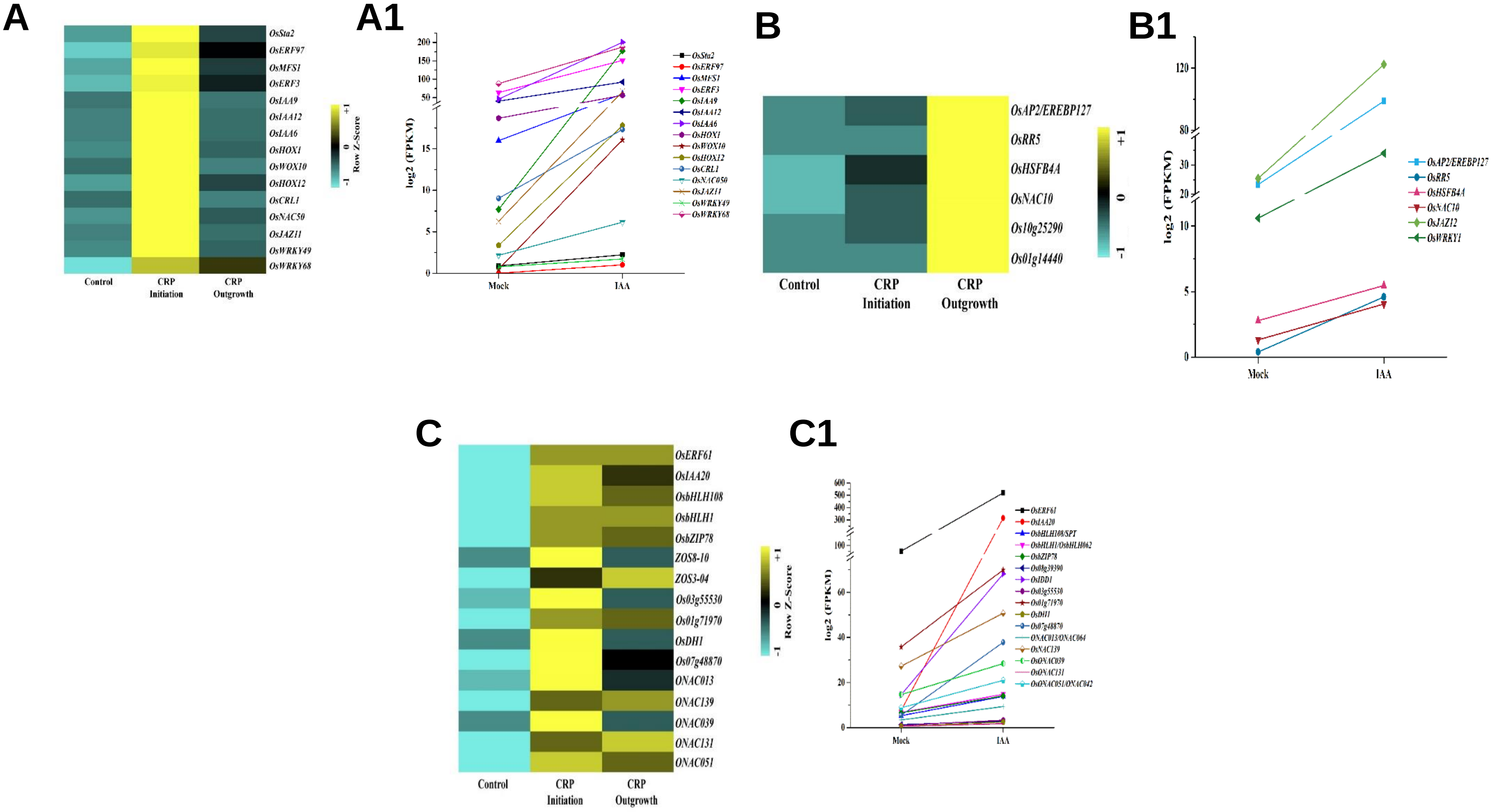


**Supplemental Figure 5:** Auxin responsiveness of differentially expressed transcriptional regulators in developing CRP. (A-C1) Expression pattern and auxin induction of TFs, activated exclusively during CRP initiation (A and A1), specifically during CRP outgrowth (B and B1), and in both the stages (C and C1). Heatmaps in (A), (B), and (C) are derived from LCM-seq data whereas plots in (A1), (B1), and (C1) are derived from auxin-responsive RNA sequencing data published by Neogy et al., (2019).

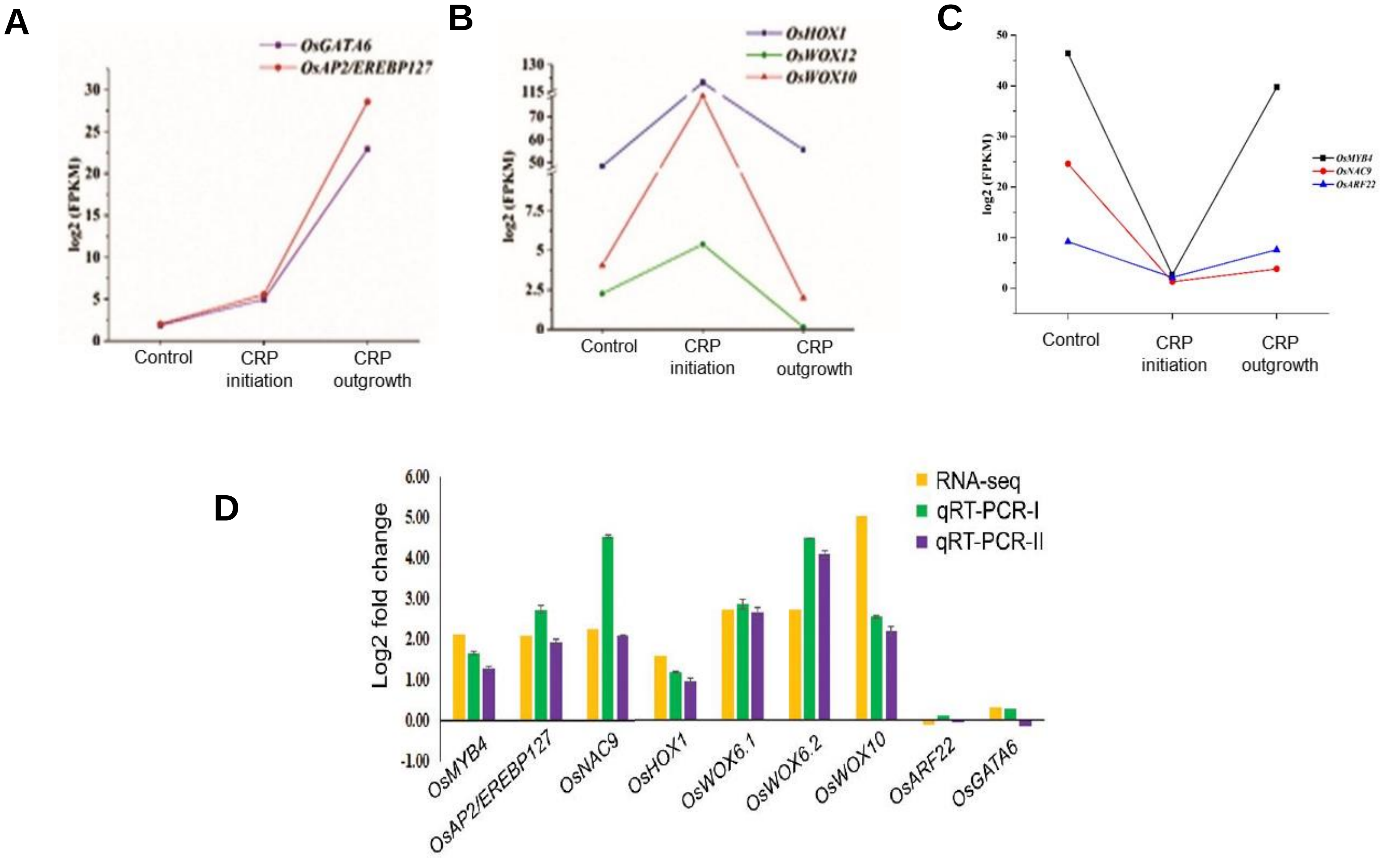


**Supplemental Figure 6:** Selected auxin-responsive CRP-expressed TFs for validation (A-C) Expression pattern of TFs during CRP development from LCM-seq data (D) Validation of these TFs for their auxin responsiveness by qRT-PCR analysis.

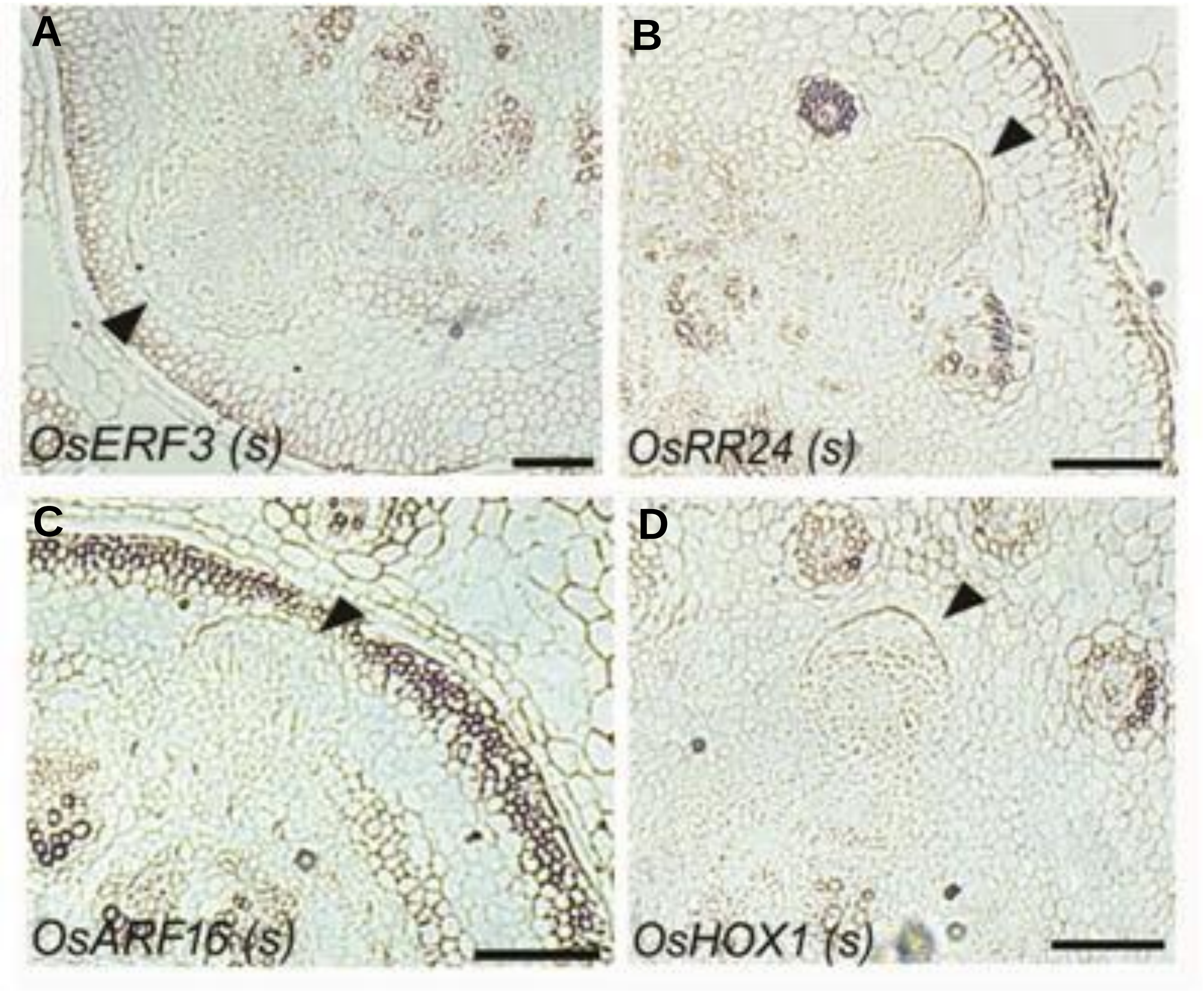


**Supplemental Figure 7:** (A-D) RNA *in situ* hybridization using sense riboprobes of *OsERF3* (A), *OsRR24* (B), *OsARF16* (C), and *OsHOX1* (D), on cross sections of stem base of wild-type plant. Bars= 100 µm in (A)-(D).

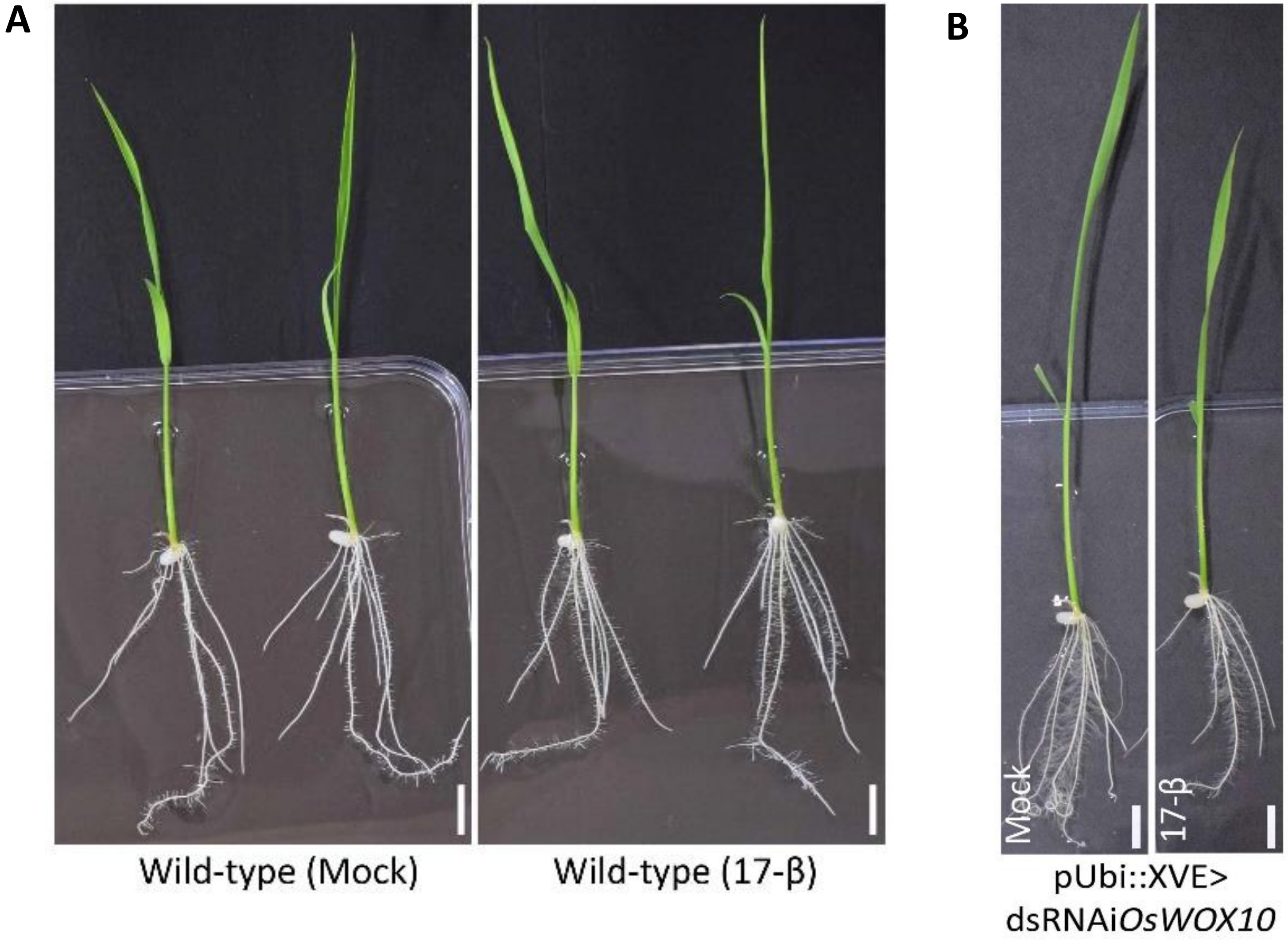


**Supplemental Figure 8:** Effects of 17-β estradiol treatment in wild-type and *OsWOX10* down-regulation. (A, B) Plant morphology of wild-type (A) and pUbi::XVE>dsRNAi*OsWOX10* lines upon mock and 17-β estradiol treatment. No significant effect was observed in wild-type plants. Bars= 1cm in (A) and (B).

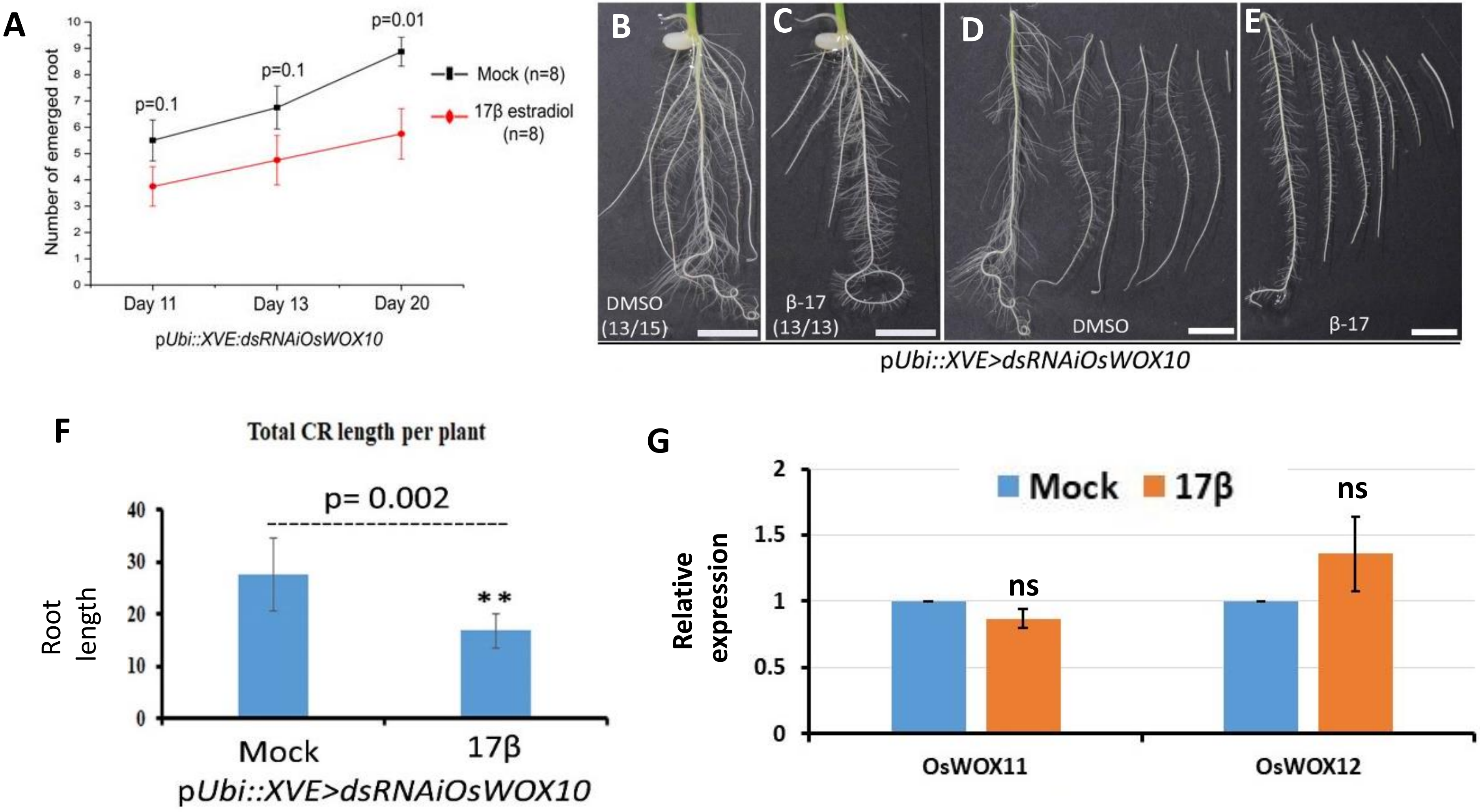


**Supplemental Figure 9:** Effects of *OsWOX10* down-regulation. (A) Root numbers are significantly reduced on 20 days after root induction of regenerating pUbi::XVE>dsRNAi*OsWOX10* lines upon estradiol treatment in the rooting media. (B-F) Plant morphology and root architecture when *OsWOX10* is down-regulated upon 17-β estradiol treatment in pUbi::XVE>dsRNAi*OsWOX10* lines in T1 generation. (G) Expression level of related WOX-genes, *OsWOX11* and *OsWOX12* upon *OsWOX10* down-regulation during CR development in T1 generation (ns, not significant; $p>0.05$; two-sample t-test). Bars= 1cm in (B)-(E).

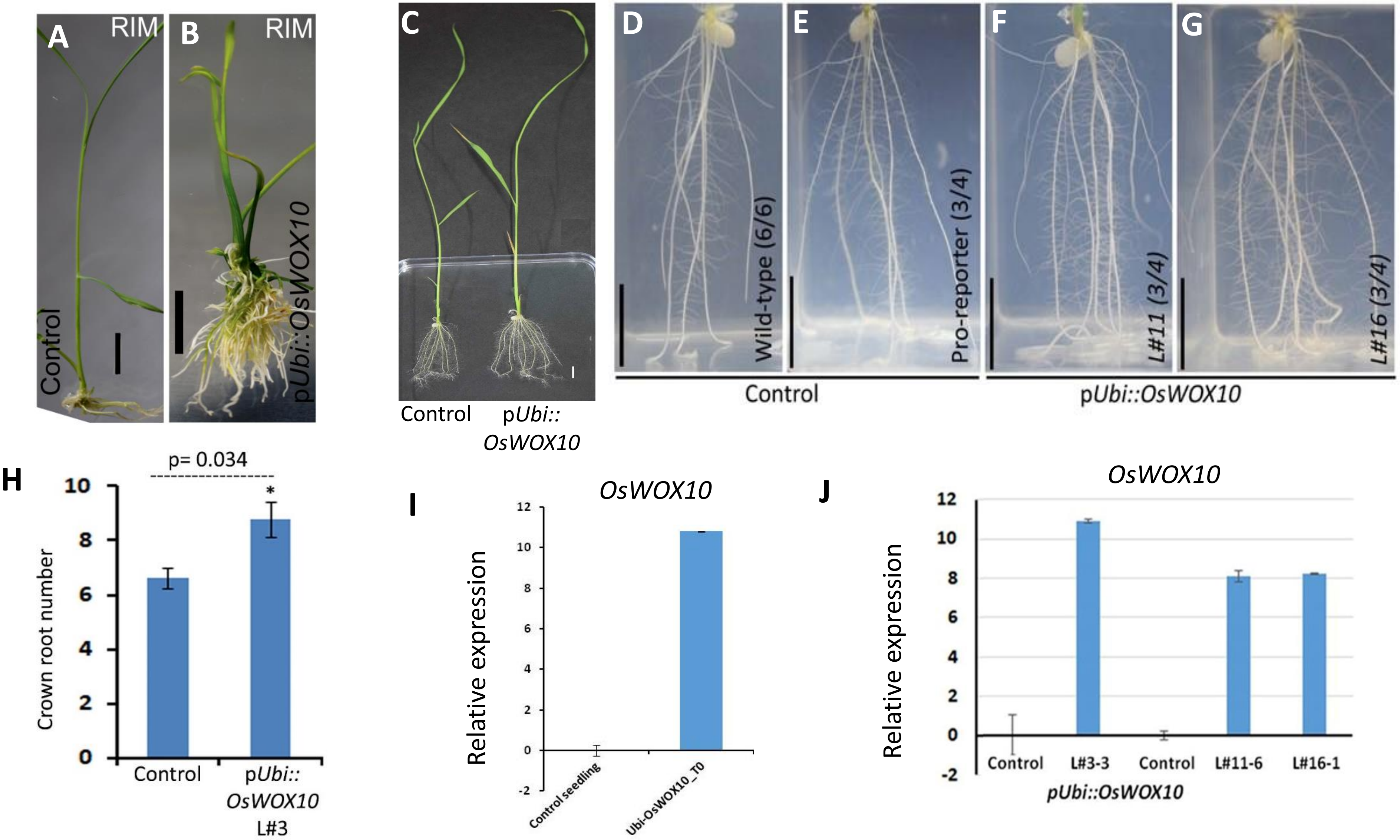


**Supplemental Figure 10:** Over-expression phenotypes of *OsWOX10*. (A, B) Extensive adventitious roots were seen in regenerating pUbi-OsWOX10 lines in root induction media (RIM). (C-G) Plant morphology and root architecture upon ectopic over-expression of *OsWOX10* during rice crown root development in T1 generation. (H) Crown root number is significantly increased in pUbi::*OsWOX10* line. (I, J) Over-expression of *OsWOX10* in p*Ubi::OsWOX10* line during plant regeneration (I) and CR development (J) by qRT-PCR analysis. Bars= 1cm in (A)-(G).

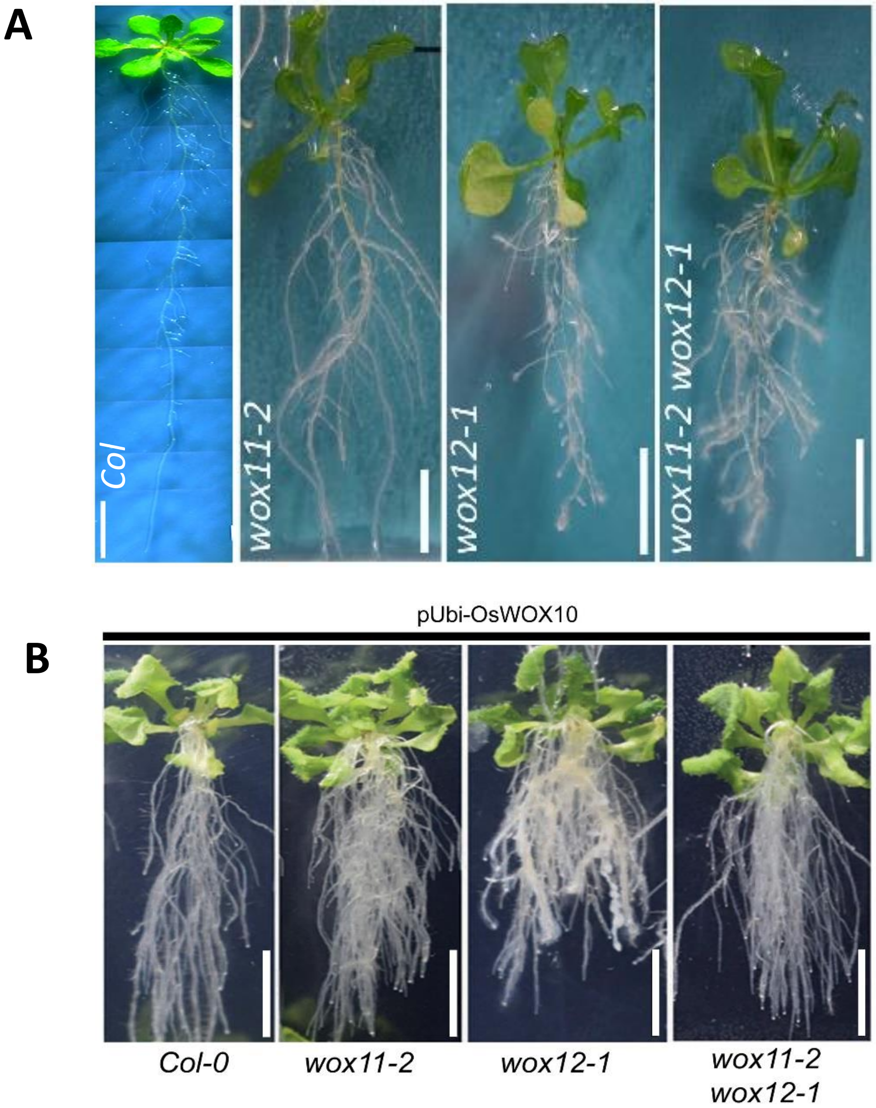


**Supplemental Figure 11:** Conserved function of *OsWOX10* in promoting adventitious root formation in *Arabidopsis*. (A, B) Effects of over-expression in three week old plants, non-transformed wild-type (*col*), *wox11-2*, *wox12-1*, and *wox11-2 wox12-1* mutants (A) and extensive root formation upon overexpression of *OsWOX10* in these backgrounds (B). Bars= 1cm in (A) and (B).

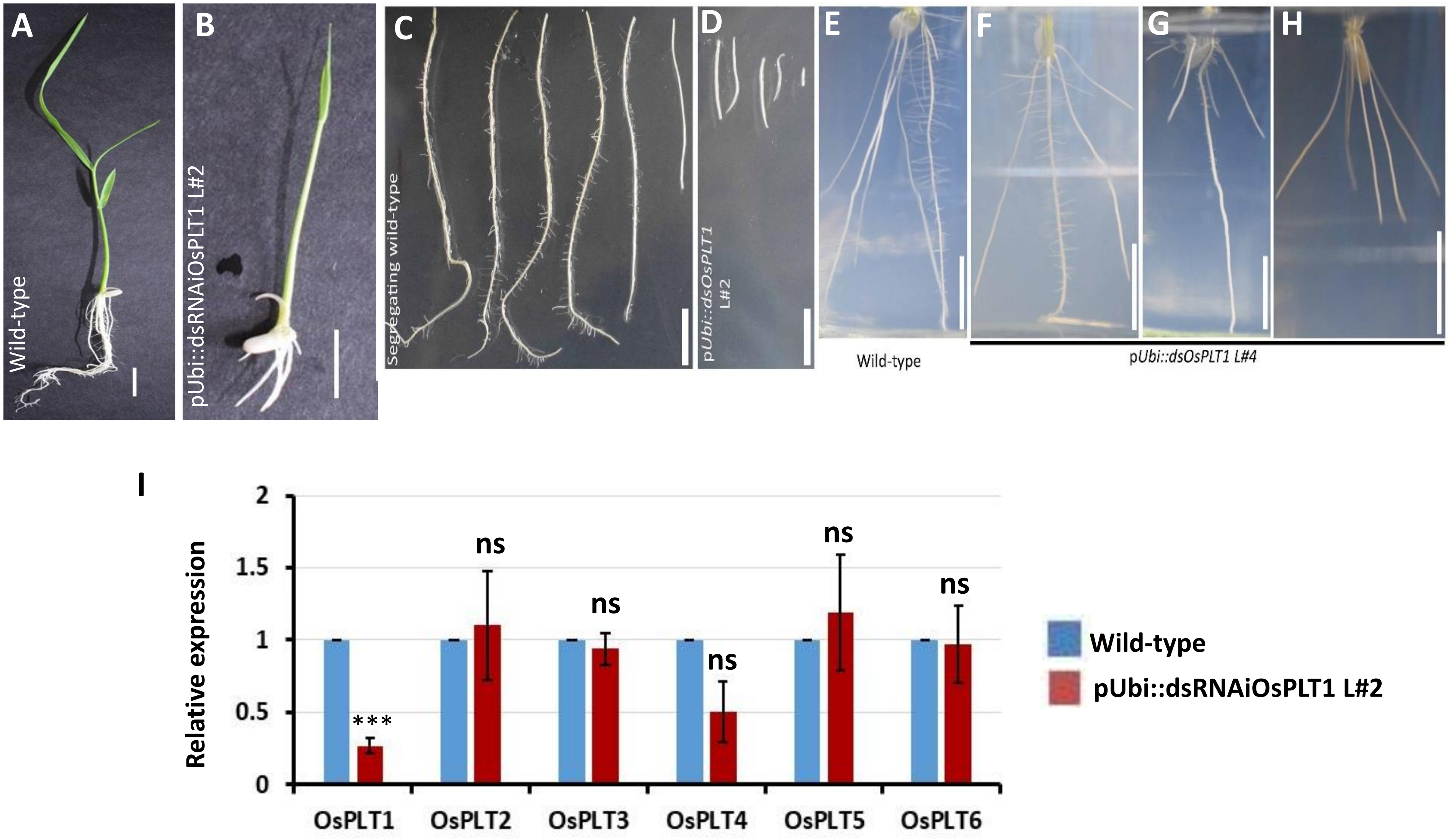


**Supplemental Figure 12:** Functional studies of *OsPLT1* during root development. (A, B) Morphology of wild-type (A) and pUbi::dsRNAi*OsPLT1* L#2 (B) plants. (C, D) Lateral root phenotype of pUbi::dsRNAi*OsPLT1* L#2. (E-H) Root architecture phenotypes pUbi::dsRNAi*OsPLT1* L#4. (I) Expression level of PLT genes, in pUbi::dsRNAi*OsPLT1* L#2 (ns, not significant; $p>0.05$; ***$p<0.001$; two-sample t-test). Bars= 1cm in (A)-(H).

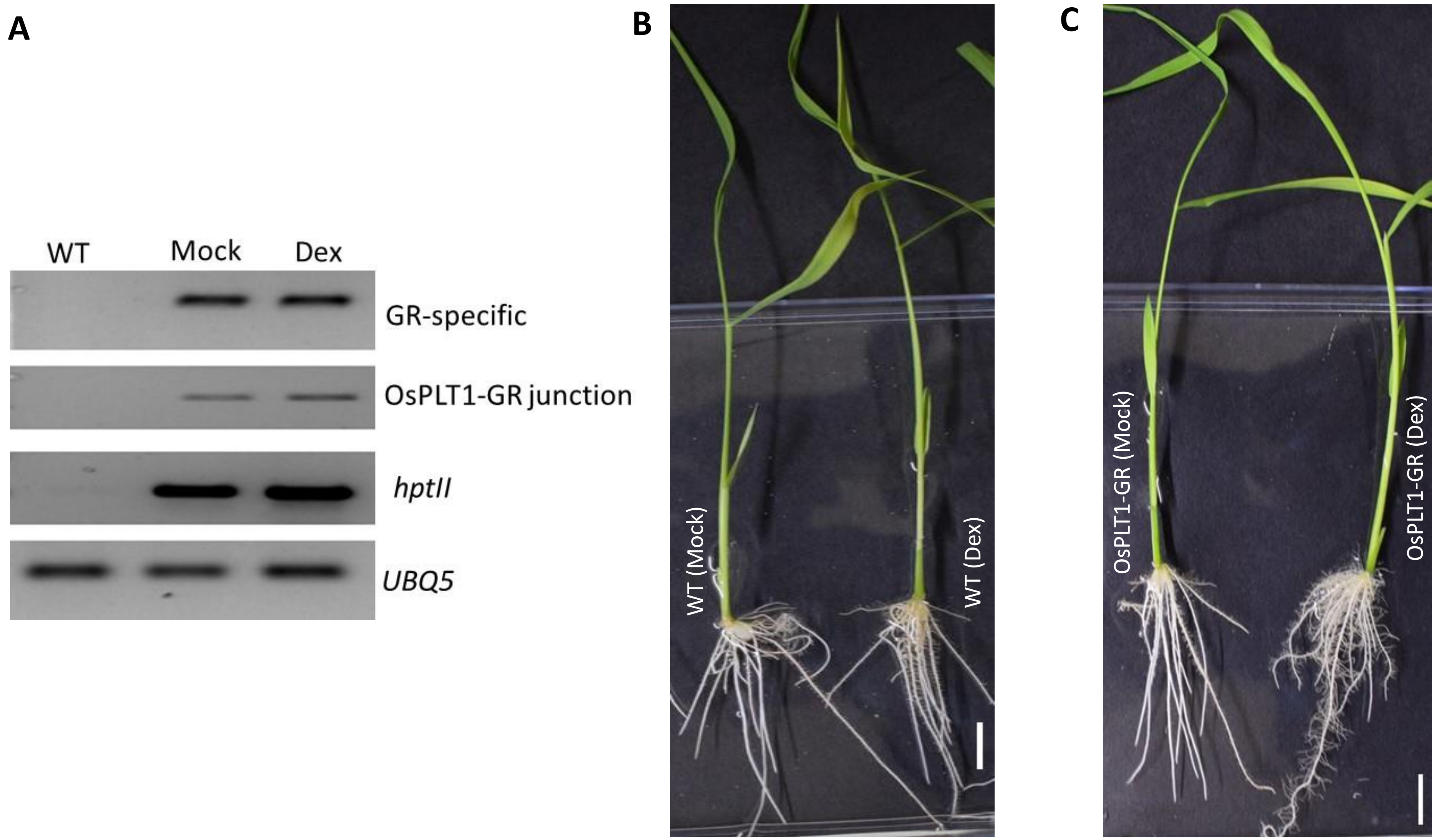


**Supplemental Figure 13:** Molecular and phenotypic characterization of *OsPLT1* over-expression lines. (A) RT-PCR analysis of *OsPLT1-GR* lines showing expression of *OsPLT1-GR* fusion transcripts. (B, C) Plant morphology of wild-type (B) and pUbi::*OsPLT1-GR* L#6 (C) upon dexamethasone treatment. No significant effect was seen on the gross morphology of wild-type plants upon dexamethasone treatment. Bars= 1cm in (B) and (C).

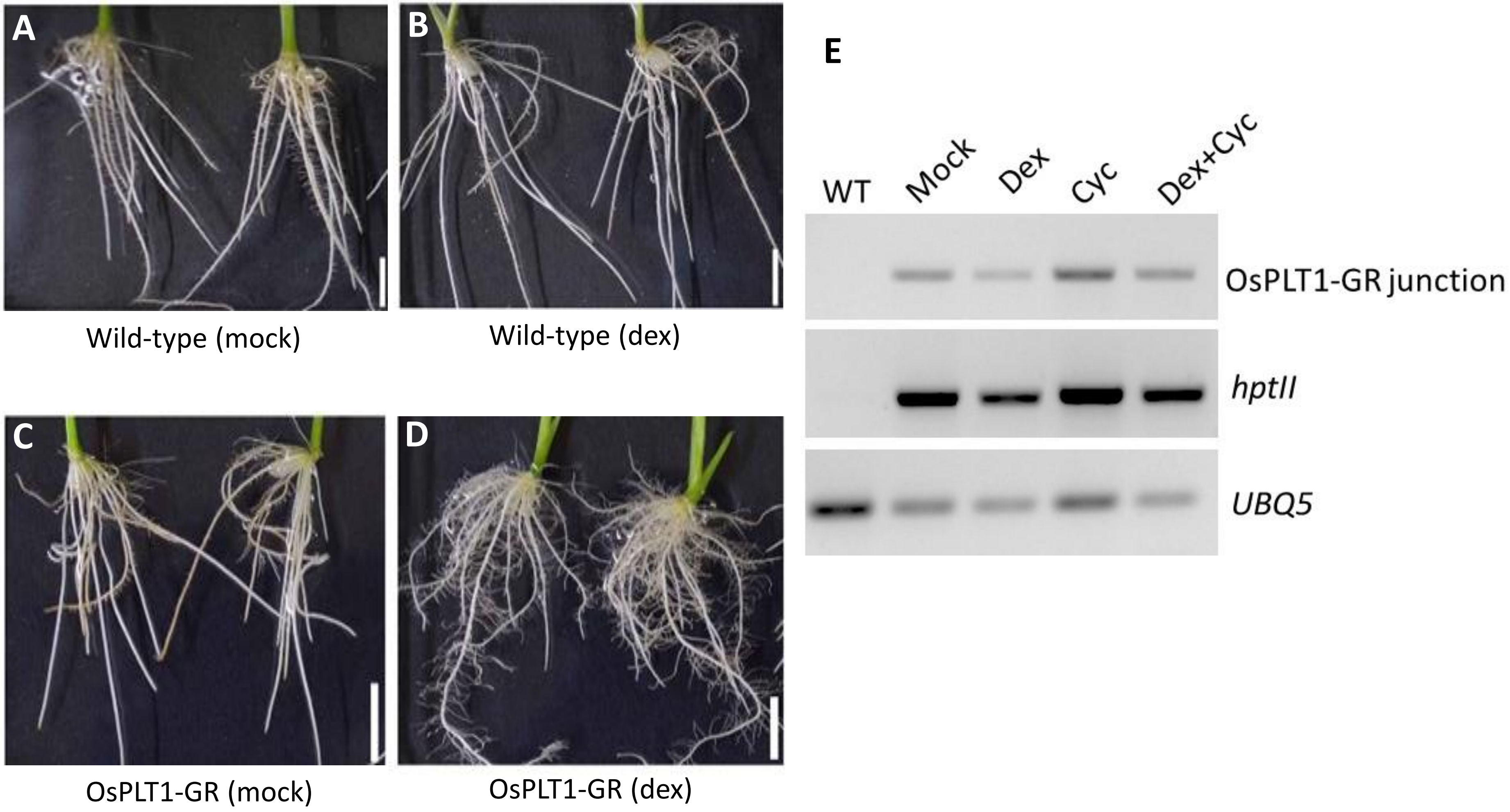


**Supplemental Figure 14:** Root architecture phenotypes of *OsPLT1* over-expression lines. (A-D) Root architecture was altered upon dexamethasone treatment in pUbi::*OsPLT1-GR* plants (D), as compared to mock treated plants (C) but no effect was seen on the root architecture of wild-type plants upon dexamethasone treatment (B) as compared to mock treated plants (A). (E) RT-PCR analysis of *OsPLT1-GR* fusion transcripts in the plants treated with Mock, dexamethasone alone (Dex), cycloheximide alone (Cyc), and dex and cyc together in combination. Bars= 1cm in (A)-(D).
